# Ribosomal S6 kinase 1 regulates ‘inflammaging’ via the senescence secretome

**DOI:** 10.1101/2023.10.17.562098

**Authors:** Suchira Gallage, Elaine E. Irvine, Silvia M.A. Pedroni, Jose Efren Barragan Avila, Sanjay Khadayate, Joaquim Pombo, Sharon Brookes, Danijela Heide, Gopuraja Dharmalingham, Agharul I. Choudhury, Nicolás Herranz, Santiago Vernia, Mathias Heikenwalder, Jesús Gil, Dominic J. Withers

## Abstract

Inhibition of the nutrient-responsive mTOR (mammalian target of rapamycin) signalling pathway including the key downstream effector S6 kinase 1 (S6K1) extends lifespan and improves healthspan in mice. However, the underlying mechanisms contributing to the broad range of age-related benefits observed with loss of S6K1 signalling are unclear. Cellular senescence is a stable growth arrest accompanied by an inflammatory phenotype (termed the senescence-associated secretory phenotype, or SASP). While both cellular senescence and SASP-mediated chronic inflammation contribute to age-related pathology, the specific role of S6K1 signalling in these processes has not been determined. Here, focussing on mouse liver, a key target tissue for the beneficial metabolic effects of loss of S6K1 signalling, we show that S6K1 deletion does not reduce senescence but ameliorates inflammation and immune cell infiltration in aged livers. Using human and mouse models of senescence, we demonstrated that reduced inflammation is a liver-intrinsic effect associated with S6K deletion. Furthermore, gene expression analysis suggested that downregulated cGAS/STING and IRF3 activation might mediate the impaired SASP observed upon S6K deletion. Using a hepatic oncogene induced senescence model, we showed *in viv*o that *S6K1* deletion results in reduced IRF3 activation, impaired production of cytokines such as IL1ý and reduced immune infiltration. Overall, deletion of S6K reduces inflammation in the liver suggesting that suppression of the inflammatory SASP by loss of S6K could contribute to explain the beneficial effects of inhibiting this pathway on healthspan and lifespan.

## INTRODUCTION

The mammalian Target of Rapamycin (mTOR) pathway plays a key role in integrating hormone and nutrient signalling and stress responses with both cellular and organismal growth and metabolism ^1^. Furthermore, mTOR signalling plays an evolutionarily conserved role in regulating longevity and healthspan ^2, 3^. For example, pharmacological inhibition of mTOR by rapamycin extends lifespan in yeast ^4^, flies ^5^ and mice ^6^. A key effector of mTOR signalling is ribosomal protein S6 kinase 1 (S6K1), which plays several roles regulating the translational machinery, cellular energy levels and has feedback effects on insulin signalling ^7, 8^. S6K1 itself has been shown to regulate ageing and different age-related processes ^9, 10^. Deletion of S6K1 extends lifespan and healthspan in mice, and also regulates longevity in flies and worms ^9, 11^.

Mice lacking S6K1 display beneficial metabolic effects including reduced adipose mass, resistance to the consequences of high fat diet feeding and increased insulin sensitivity ^12, 13^, a constellation of phenotypes that align with the effects of calorie restriction (CR) a conserved longevity mechanism ^14^. Different molecular mechanisms have been proposed to explain these effects. For example, loss of S6K1 leads to upregulation of the activity of AMP kinase a key regulator of cellular energy homeostasis ^15^, thus mimicking the effects of CR and motivating the use of metformin as a potential geroprotective drug ^16^. S6K1 also phosphorylates the glutamyl-prolyl-tRNA synthetase (EPRS), which in turn is involved in regulating adiposity and adipose tissue metabolism and this may underlie the beneficial metabolic phenotypes observed in S6K1 null mice ^17^. Despite these insights, a definitive answer to what are the cellular and molecular mechanisms behind the broad-ranging effects associated with abrogation of S6K signalling is unclear.

Cellular senescence is a stress response that limits the replication of old, damaged and cancerous cells ^18^. Senescent cells stably exit cell cycle and undergo multiple phenotypic changes, including the production of a pro-inflammatory secretome known as the senescence-associated secretory phenotype (SASP) ^19^. Cellular senescence is a hallmark of ageing ^20^: senescent cells not only accumulate during ageing ^21^ but contribute to ageing and age-related diseases ^22^.

Interestingly mTOR influences different phenotypes associated with senescence ^14^. Inhibition of mTOR prevents senescence by interfering with the establishment of an irreversible growth arrest ^23, 24^. On the other hand, treatment of already senescent cells with rapamycin inhibits the inflammatory SASP ^25, 26^. Mechanistically, 4EBP has been implicated in mTOR-mediated SASP regulation ^25, 26^, but a role for S6K signalling has not been investigated. Moreover, the S6K-STING interaction has been shown to regulate immune responses ^27^. Since the cGAS/STING pathway is central to regulate the SASP ^28, 29^, it is tempting to hypothesise whether S6K could regulate the inflammatory SASP.

Given our incomplete understanding of the mechanisms by which loss of S6K1 signalling benefits ageing and age-related pathologies, we undertook a series of studies in long-lived *S6K1^-/-^*mice and other genetic and pharmacological models of S6K inhibition. We explored the role of senescence, the SASP and inflammation in the liver, as this organ displays several age-related changes (including an increased inflammatory profile, ^30, 31^) and shows beneficial metabolic phenotypes in mice lacking S6K1 ^32, 33^. In these studies, we find that loss of S6K1 attenuates age-related liver pathology, does not influence senescence but reduced liver inflammation via effects on the proinflammatory SASP and immune surveillance. Thus, S6K signalling plays a key role in age-related inflammation (inflammaging ^34^) and targeting this pathway may be a strategy for treating the diseases of ageing.

## RESULTS

### *S6K1* deletion attenuates age-related liver pathology

To investigate the role of S6K1 in age-related liver pathology, particularly in the regulation of senescence and the SASP, we compared the livers of old (600 day old) *S6K1* WT and KO female mice (Fig. 1a-b). To this end, we established two cohorts of mice. As previously described, *S6K1* KO mice were smaller than age-matched *S6K1* WT littermates (Extended Data Fig. 1a-b) and displayed significantly reduced epididymal white adipose tissue (eWAT) and liver mass (Extended data Fig. 1c and Fig. 1c).

**Figure 1.**
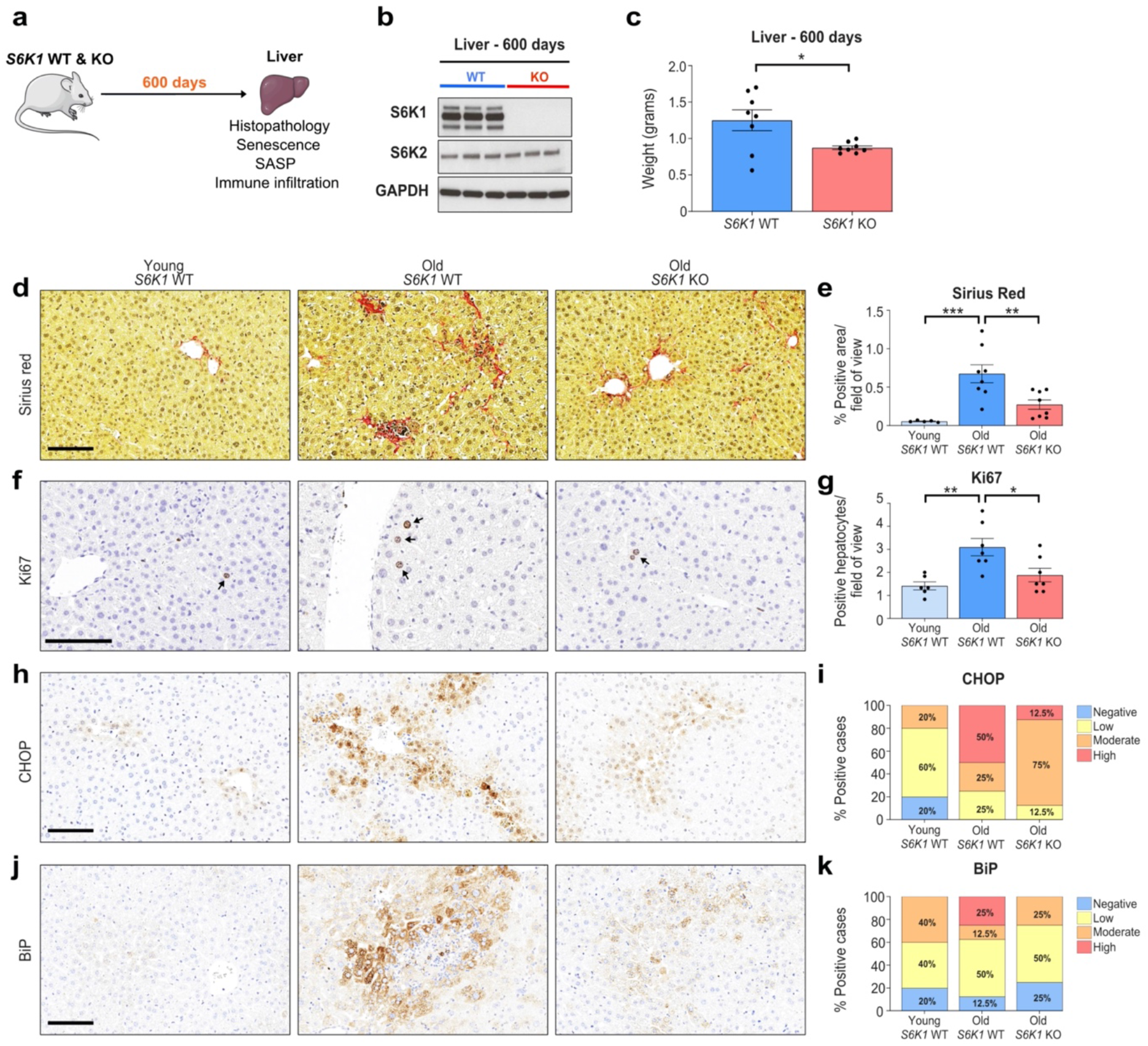
S6K1 deletion attenuates age-related liver pathology. **a.** Experimental scheme. *S6K1* WT and KO mice were aged for 600 days to assess senescence. **b.** Representative immunoblot analysis of S6K1, S6K2 and GAPDH protein expression in whole liver lysates of 600-day-old *S6K1* WT (left; n=3) and KO (right; n=3) mice. S6K1 and GAPDH (loading control) were run on the same blot. S6K2 was run on a separate blot. **c.** Liver weight (grams) at 600 days from *S6K1* WT (n=8) and KO (n=8) mice. **d, e.** Sirius red staining **(left)** and quantification **(right)** in livers in young *S6K1* WT (90 days; n=5), old *S6K1* WT (600 days; n=8) and old *S6K1* KO (600 days; n=8) mice. **f, g.** Ki67 staining **(left)** and quantification **(right)** in livers in young *S6K1* WT (90 days; n=6), old *S6K1* WT (600 days; n=7) and old *S6K1* KO (600 days; n=7) mice. **h,i.** CHOP staining **(left)** and quantification **(right)** in livers in young *S6K1* WT (90 days; n=5), old *S6K1* WT (600 days; n=8) and old *S6K1* KO (600 days; n=8) mice. **j, k.** BiP staining **(left)** and quantification **(right)** in livers in young *S6K1* WT (90 days; n=5), old *S6K1* WT (600 days; n=8) and old *S6K1* KO (600 days; n=8) mice. Data are expressed as mean ± SEM. Statistical significance was calculated using either Student’s t-test (c) or one-way analysis of variance with Tukey’s multiple comparison test (e,g). (* P < 0.05, ** P < 0.01, *** P < 0.001). *n* denotes individual mice. Scale bar, 100 μm (d,h,j) or 50 μm (f). n.s: non-significant. WT: wild type. KO: knockout.

Senescent cells contribute to age-related liver pathology including hepatic steatosis ^35^ and inflammation ^31^ as well as liver fibrosis ^36, 37^. Consistent with previous evidence of preserved organ homeostasis and function at old age, *S6K1* KO mice showed improved liver pathology (Fig. 1d-k). Sirius Red staining of liver sections showed that old *S6K1* WT mice displayed increased levels of fibrosis than their younger counterparts, whereas fibrosis was significantly lower in old *S6K1* KO littermates (Fig. 1d-e). An increase in the numbers of Ki67+ hepatocytes observed in the livers of old *S6K1* WT mice, indicative of compensatory proliferation in response to age-related liver damage, was reduced in the livers of old *S6K1* KO littermates (Fig. 1f-g). Markers of ER stress such as CHOP and BiP, that reflect liver damage, were also found elevated in the livers of old *S6K1* WT mice, whereas this was attenuated in the livers of old *S6K1* KO littermates (Fig. 1h-k). Overall, these data confirm that old *S6K1* KO mice display increased liver fitness as reflected by an amelioration of age-related liver pathology.

### S6K1 status affects inflammation but not senescence in the livers of old mice

Senescent cells accumulate during ageing ^21^ and contribute to age-related tissue dysfunction through the production of the SASP ^38^.

We hypothesised that regulation of senescence by S6K1 may explain, at least in part, the beneficial effects that *S6K1* loss has on age-related liver pathology. However, markers of senescence such as the Cdkn2a transcripts encoding for p16^Ink4a^ (Fig. 2a) and p19^Arf^ (Fig. 2b) were upregulated in old mice irrespective of the genotype, suggesting that S6K1 status did not affect the senescence response *per se*. Moreover, gene set-enrichment analysis (GSEA) performed on RNA-Seq of liver samples (Fig. 2c) unveiled senescence-related signatures were upregulated in old animals, irrespective of S6K1 status (Fig. 2d), further suggesting that S6K1 loss did not affect senescence induction.

**Figure 2.**
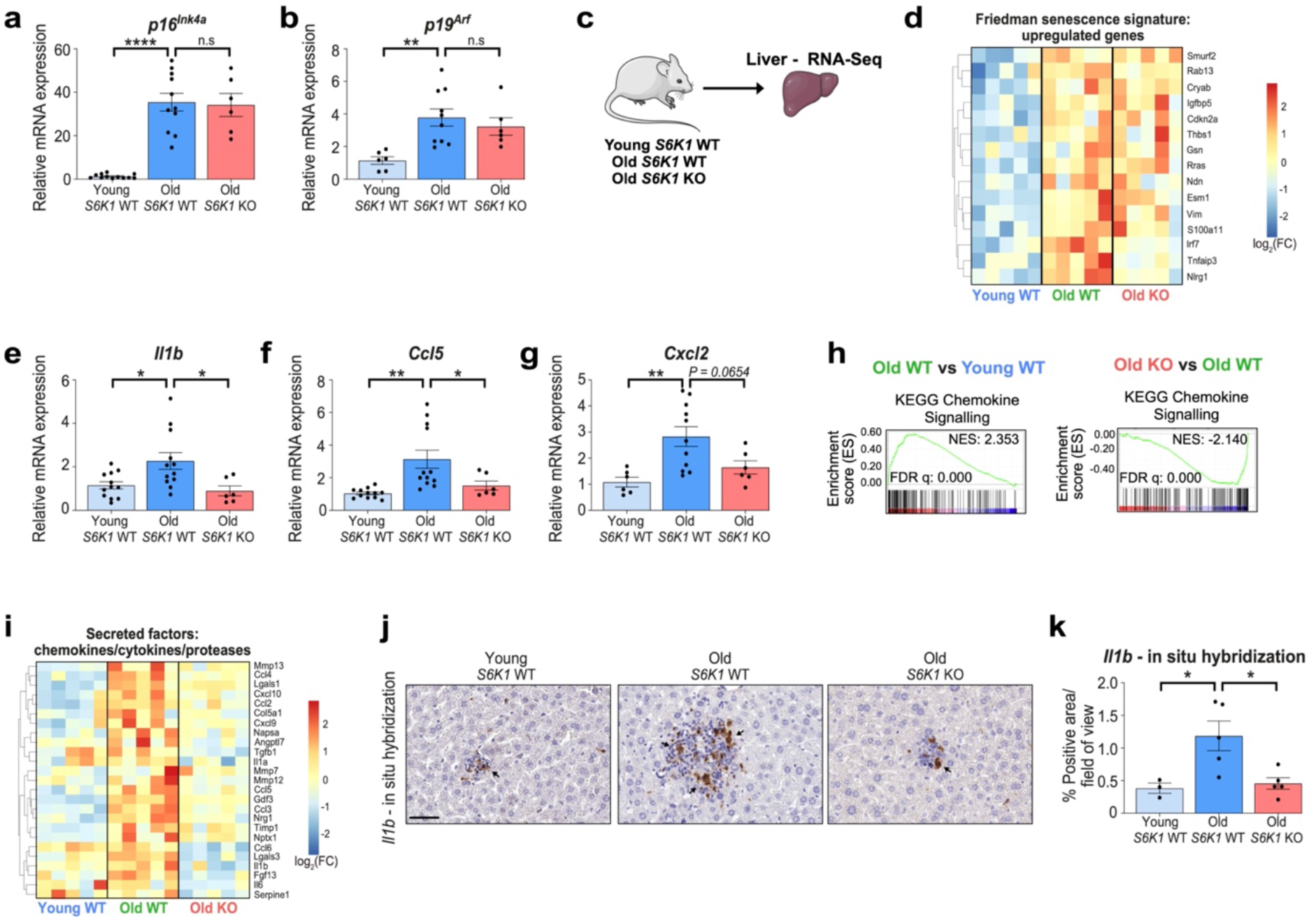
S6K1 status affects inflammation but not senescence in the livers of old mice. **a,b.** Relative mRNA expression for *p16^Ink^*^4a^ and *p19^Arf^* assessed by RT-qPCR from whole liver lysates of young *S6K1* WT (90 days; n=12 *p16^Ink^*^4a^ and n=6 *p19^Arf^*), old *S6K1* WT (600 days; n=11 *p16^Ink^*^4a^ and n=10 *p19^Arf^*) and old *S6K1* KO (600 days; n=6 for *p16^Ink^*^4^ and *p19^Arf^*) mice. mRNA expression was normalized to Rps14 housekeeping gene. **c.** Experimental scheme. Bulk RNA-sequencing of whole liver lysates from young *S6K1* WT (90 days), old *S6K1* WT (600 days) and old *S6K1* KO (600 days) mice. **d.** Heatmap depicting the expression of key regulated genes in the ‘Friedman senescence signature’ in young *S6K1* WT (90 days; n=5), old *S6K1* WT (600 days; n=5) and old *S6K1* KO (600 days; n=5) mice. **e-g.** Relative mRNA expression for *Il1b, Ccl5 and Cxcl2* assessed by RT-qPCR from whole liver lysates of young *S6K1* WT (90 days; n=12 for *Il1b* and *Ccl5;* n=6 for *Cxcl2*), old *S6K1* WT (600 days; n=12 for *Il1b* and *Ccl5;* n=11 for *Cxcl2*) and old *S6K1* KO (600 days; n=6 for *Il1b, Ccl5 and Cxcl2*) mice. mRNA expression was normalized to Rps14 housekeeping gene. **h.** Gene-set enrichment analysis (GSEA) for ‘KEGG Chemokine Signalling’ of young *S6K1* WT (90 days), old *S6K1* WT (600 days) and old *S6K1* KO (600 days) mice from whole liver lysates. **i.** Heatmap depicting the expression of key chemokines, cytokines and proteases in young *S6K1* WT (90 days; n=5), old *S6K1* WT (600 days; n=5) and old *S6K1* KO (600 days; n=5) mice. **j.k.** *In situ* hybridization for *Il1b* mRNA **(left)** and quantification **(right)** in young *S6K1* WT (90 days; n=3), old *S6K1* WT (600 days; n=5) and old *S6K1* KO (600 days; n=5) mice. Data are expressed as mean ± SEM. Statistical significance was calculated using one-way analysis of variance with Tukey’s multiple comparison test. (* P < 0.05, ** P < 0.01, **** P < 0.0001). *n* denotes individual mice. Scale bar, 100 μm (d,f,h,j) or 50 μm (l). n.s: non-significant. WT: wild type. KO: knockout. NES: normalized enrichment score. FDR: false discovery rate.

Senescent cells produce a complex mix of immunomodulatory cytokines referred to as the senescence-associated secretory phenotype (SASP) ^19^. In this regard, the expression of several immunomodulatory cytokines in the liver, including *Il1b*, *Ccl5* and *Cxcl2* (Fig. 2e-g) were elevated during ageing in *S6K1* WT mice but were downregulated in old *S6K1* KO animals. Furthermore, GSEA showed that a chemokine signalling signature was elevated in old *S6K1* WT mice when compared with young control animals, and this was also downregulated in old *S6K1* KO littermates (Fig. 2h). Heatmaps confirmed that the expression of multiple cytokines increased with age in the livers of *S6K1* WT while remaining at lower levels in *S6K1* KO mice (Fig. 2i). Finally, RNA in situ hybdrdization of liver sections showed that *Il1b* was elevated in livers of old *S6K1* WT mice, when compared to livers of old *S6K1* KO mice (Fig. 2j-k). These results suggest that *S6K1* loss does not affect senescence but rather impairs the production of a subset of inflammatory cytokines during ageing.

### *S6K1* deletion prevents inflammaging in livers

The accumulation of senescent cells has been associated with chronic inflammation and persistent immune-cell infiltrates during ageing, often referred to as inflammaging ^34, 38^. Given that the livers of old *S6K1* WT mice displayed lower expression of inflammatory, immunomodulatory cytokines, we speculated that this might result in decreased levels of chronic immune infiltration in old mice. GSEA showed that an immune response signature was significantly upregulated in livers of old *S6K1* WT mice when compared to either young mice or old *S6K1* KO mice (Figure 3a). Moreover, H&E staining of liver sections showed increased immune infiltration in old *S6K1* WT mice that was less pronounced in age-matched *S6K1* KO littermates (Fig. 3b). To study this in more detail, we used quantitative immunohistochemistry on whole liver sections for various immune cell markers. We have previously observed an increase in myeloid and lymphoid infiltrates in the liver of aged mice ^31^. In agreement with those results, an increase in myeloid (major histocompatibility complex II (MHC-II+, CD68+, F4/80+) and lymphoid (CD3+ and B220+) infiltrates was observed in old *S6K1* WT mice. Interestingly, age-matched *S6K1* KO littermates display reduced infiltration of myeloid cells (MHCII+, CD68+ and F4/80+ cells; Fig. 3c-h), T cells (CD3+ or CD4+ cells; Fig. 3i-j and Extended Data Fig. 2a-b) and B cells (B220+ cells; Fig. 3k-l). We did not observe significant differences in the presence of platelet infiltration (Extended Data Fig. 2c-d), suggesting that the effect was specific.

**Figure 3.**
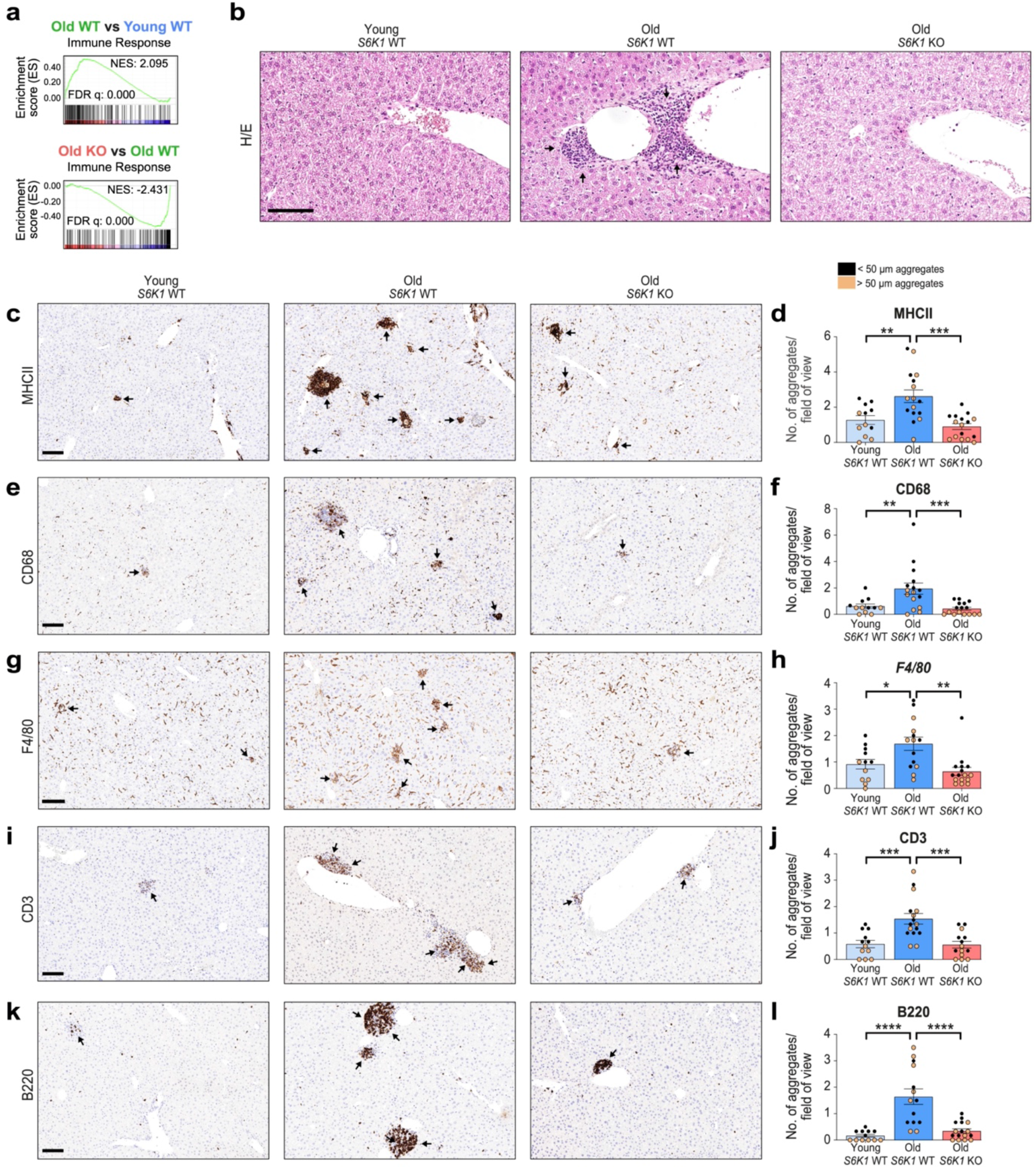
S6K1 deletion prevents inflammaging in livers. **a. h.** Gene-set enrichment analysis (GSEA) for ‘Immune Response’ of young *S6K1* WT (90 days), old *S6K1* WT (600 days) and old *S6K1* KO (600 days) mice from whole liver lysates. **b.** Haematoxylin and eosin (H/E) staining of livers from mice of the indicated genotypes. **c,d.** MHCII staining for antigen presenting cells **(left)** and quantification **(right)** of livers from young *S6K1* WT (90 days; n=6), old *S6K1* WT (600 days; n=8) and old *S6K1* KO (600 days; n=8) mice. **e,f.** CD68 staining for monocytes and macrophages **(left)** and quantification **(right)** of livers from young *S6K1* WT (90 days; n=6), old *S6K1* WT (600 days; n=8) and old *S6K1* KO (600 days; n=8) mice. **g,h.** F4/80 staining for resident Kupffer cells **(left)** and quantification **(right)** of livers from young *S6K1* WT (90 days; n=6), old *S6K1* WT (600 days; n=7) and old *S6K1* KO (600 days; n=8) mice. **i, j.** CD3 staining for T-cells **(left)** and quantification **(right)** of livers from young *S6K1* WT (90 days; n=6), old *S6K1* WT (600 days; n=8) and old *S6K1* KO (600 days; n=7) mice. **k, l.** B220 staining for B-cells **(left)** and quantification **(right)** of livers from young *S6K1* WT (90 days; n=6), old *S6K1* WT (600 days; n=7) and old *S6K1* KO (600 days; n=8) mice. Data are expressed as mean ± SEM. Statistical significance was calculated using one-way analysis of variance with Tukey’s multiple comparison test. (* P < 0.05, ** P < 0.01, *** P < 0.001, **** P < 0.0001). *n* denotes individual mice. Scale bar, 100 μm. n.s: non-significant. WT: wild type. KO: knockout. NES: normalized enrichment score. FDR: false discovery rate.

There were no striking differences in immune infiltrates in the livers of young *S6K1* WT and KO mice (Extended Data Fig. 3), nor were there differences in the levels of circulating monocytes/lymphocytes in the blood taken from the respective cohorts (Extended Data Fig. 4). These results suggest that the lower infiltration of specific immune cell types observed in livers of old *S6K1* KO mice was due to a specific effect on recruitment into the liver, rather than an alteration in populations or numbers in peripheral blood. Overall, the above results suggest that while S6K1 loss does not result in reduced numbers of senescent cells in liver of old wild-type mice, livers of old *S6K1* KO mice showed less chronic inflammation and immune infiltration than old *S6K1* WT controls.

### S6K regulates the inflammatory SASP in MEFs

A possible explanation for the phenotypes observed in the livers of *S6K1* KO mice is that *S6K1* loss results in a selective inhibition of proinflammatory cytokines without affecting other senescence phenotypes. To investigate this possibility, we took advantage of mouse embryonic fibroblasts (MEFs) derived from *S6K1* KO, *S6K2* KO or *S6K1/S6K2* DKO mice and compared them to their WT counterparts (Fig 4a).

**Figure 4.**
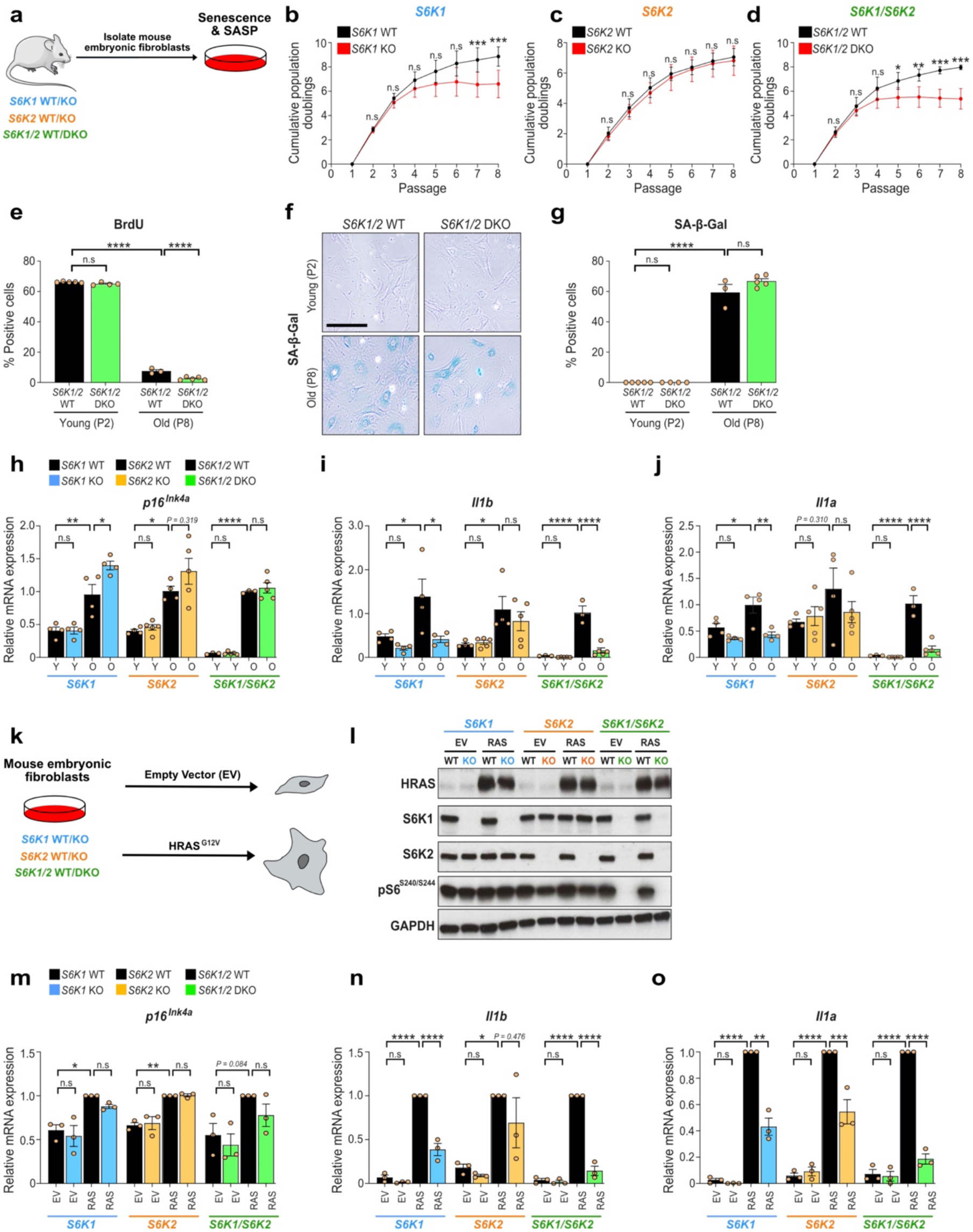
S6K1 and/or S6K2 deletion does not bypass senescence but dampens SASP induction in mouse embryonic fibroblasts. **a.** Experimental scheme. Mouse embryonic fibroblasts (MEFs) were generated from *S6K1* WT/KO, *S6K2* WT/KO and *S6K1/2* WT/DKO embryos and were assessed for replicative senescence. MEFs were generated from 3-5 independent pairs of embryos from at least 3 different mothers. **b-d.** Cumulative population doublings of *S6K1* WT (n=5) and KO (n=4) MEFs **(left)**, *S6K2* WT (n=4) and KO (n=5) MEFs **(middle)** and *S6K1/2* WT (n=3) and DKO (n=5) MEFs **(right)**. **e.** Quantification of BrdU incorporation in young (passage 2) and old (passage 8) *S6K1/2* WT (young n=5; old n=3) and DKO (young n=4; old n=5) cells. **f,g.** Representative images **(left)** and quantification **(right)** of senescence-associated beta galactosidase (SA-β-Gal) staining in young (passage 2) and old (passage 8) *S6K1/2* WT (young n=5; old n=3) and DKO (young n=4; old n=5) cells. **h-j.** Relative mRNA expression for *p16^Ink^*^4a^ **(left)**, *Il1b* **(middle)** and *Il1a* **(right)** assessed by RT-qPCR from young (passage 3) and old (passage 8) MEFs from *S6K1* WT (n=4) and KO (n=4), *S6K2* WT (n=4) and KO (n=5) as well as *S6K1/2* WT (n=3) and DKO (n=5) cells. mRNA expression was normalized to *Rps14* housekeeping gene. **k.** Experimental scheme. MEFs of the indicated genotypes were stably transduced with a retroviral vector containing the empty vector or expressing HRAS^G12V^. MEFs were generated from 3 independent pairs of embryos from 3 different mothers. **l.** Immunoblot analysis for HRAS, S6K1, S6K2, pS6^S240/S244^ and GAPDH expression. S6K1 and GAPDH (loading control) were run on the same blot. S6K2, pS6^S240/S244^ and HRAS were run on separate blots. **m-o.** Relative mRNA expression for *p16^Ink^*^4a^ **(left)**, *Il1b* **(middle)** and *Il1a* **(right)** assessed by RT-qPCR from MEFs transduced with empty vector (EV) or *HRAS^G12V^* (RAS) from *S6K1* WT (n=3) and KO (n=3), *S6K2* WT (n=3) and KO (n=3) as well as *S6K1/2* WT (n=3) and DKO (n=3) cells. mRNA expression was normalized to *Rps14* housekeeping gene. Data are expressed as mean ± SEM. n represents Statistical significance was calculated using repeated two-way analysis of variance with Sidak’s multiple comparison test (b-d) or by two-way analysis of variance with Tukey’s multiple comparison test (e,g,h-j,m-o). (* P < 0.05, ** P < 0.01, *** P < 0.001, **** P < 0.0001). *n* denotes individual MEFs generated from distinct embryos. n.s: non-significant. WT: wild type. KO: knockout. DKO: double knockout.

Loss of *S6K1* and/or *S6K2* did not abrogate the growth arrest observed during serial passages of MEFs. Indeed, *S6K1* KO and *S6K1/S6K2* DKO cells arrested even earlier than WT MEFs (Fig 4b-d). This premature arrest, observed in *S6K1* KO and *S6K1/S6K2* DKO MEFs subjected to serial passage, was not due to intrinsic differences in cell growth (Extended Data Fig 5a-f). Analysis of WT and *S6K1/S6K2* DKO MEFs confirmed that old *S6K1/S6K2* DKO MEFs underwent senescence with similar dynamics to their WT counterparts as shown in BrdU incorporation (Fig. 4e) and SA-ý-Gal cell staining (Fig. 4f-g) assays performed on late passage MEFs. Further analysis confirmed that the transcript encoding for p16^Ink4a,^ a CDKI necessary for senescence growth arrest, was similarly induced in old MEFs regardless of *S6K1* and *S6K2* deletion (Fig.4h). Nevertheless, the induction of SASP components such as the proinflammatory cytokines *Il1a* and *Il1b* observed in late passage MEFs was significantly reduced in *S6K1* KO and *S6K1/S6K2* DKO MEFs when compared to WT cells (Fig 4i-j). We next analysed whether the observed effects are conserved during oncogene-induced senescence (OIS). We, therefore, infected MEFs with an oncogenic *Ras* (HRAS^G12V^) expressing vector or its parental vector as a control (Fig. 4k-l). *Ras* expression triggered senescence, as shown by *Ink4a* induction regardless of *S6K1* and *S6K2* status (Fig. 4m). Interestingly, the induction of *Il1a* (Fig. 4n) and *Il1b* (Fig. 4o) observed during OIS was significantly reduced in *S6K1* KO and *S6K1/S6K2* DKO MEFs when compared to WT cells. The above results suggest that deletion of *S6Ks* results in impaired production of proinflammatory cytokines during senescence with combined loss of *S6K1* and *S6K2* having the strongest effect.

### S6K1/2 regulates the proinflammatory SASP in human cells

To understand if the above results can be extended to human cells, we took advantage of IMR90 ER:RAS cells (Fig. 5a), a widely used model to study OIS in human fibroblasts ^39^. Treatment with 4-hydroxytamoxifen (4OHT) activates RAS in these cells, inducing senescence and the SASP (Fig. 5b-c). To analyse the role of S6K1 and S6K2, we took advantage of two independent siRNAs targeting each gene (Extended Data Fig. 6a, b).

**Figure 5.**
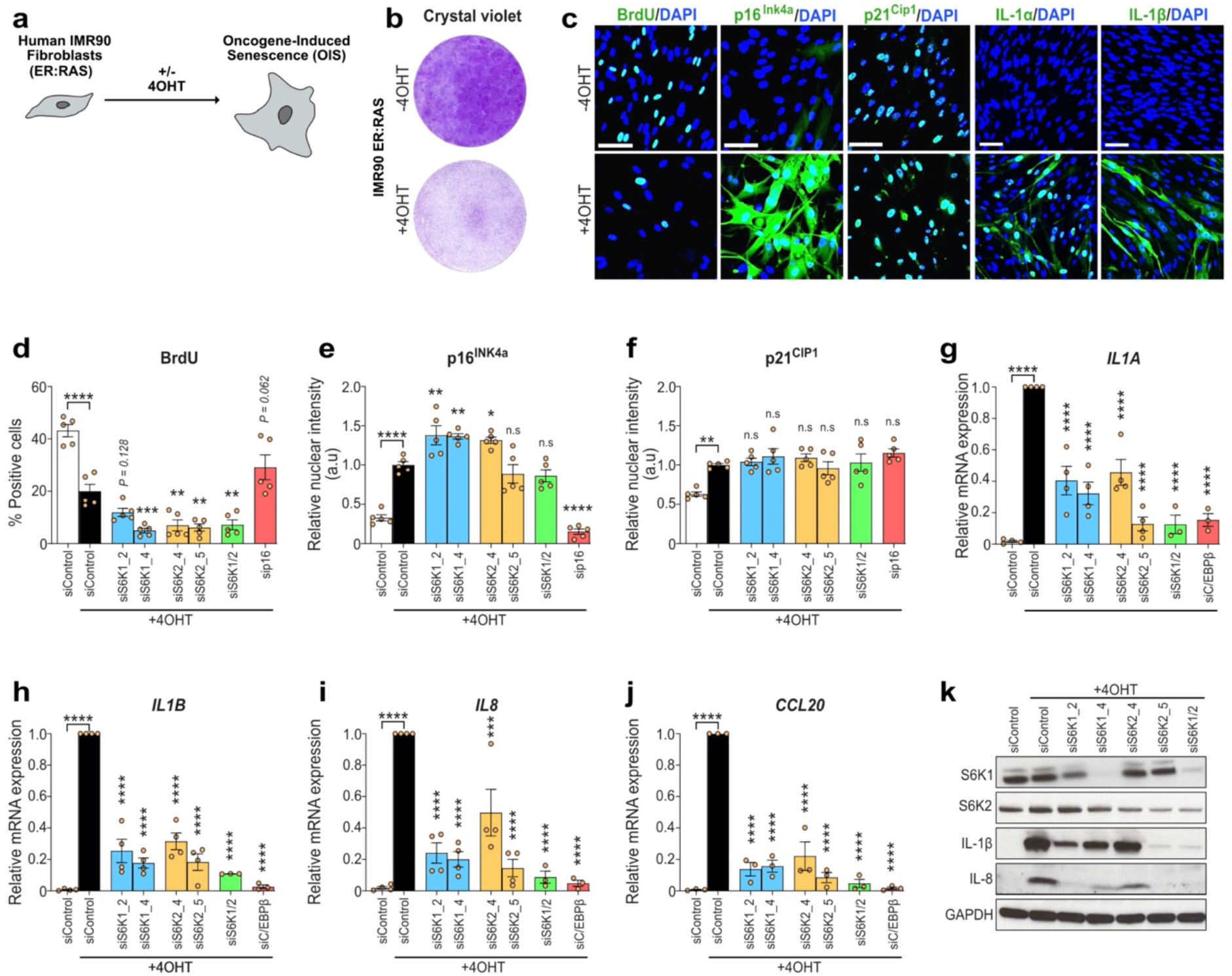
S6K1/2 regulates the SASP without affecting the growth arrest in oncogene-induced senescence in human fibroblasts. **a.** Experimental scheme. IMR90 fibroblasts were stably transduced with the pLNC-ER: RAS retroviral vector and treated with 4-hydroxytamoxifen (4OHT) for senescence induction. **b.** Cell proliferation assessed by crystal violet staining following 13 days with and without 4OHT treatment in IMR90 ER:RAS cells. **c.** Representative immunofluorescence (IF) staining of BrdU, p16^INK4A^, IL-1α, IL-1β and IL-8 following 7 days (BrdU and p16^Ink4a^) or 8 days (SASP) with or without 4OHT treatment in IMR90 ER: RAS cells. **d-f**. IMR-90 ER: RAS cells were reverse transfected with either Allstars (scrambled sequence - siControl) or the indicated siRNAs. Cells were treated with or without 4OHT on the following day to induce senescence. Quantification of immunofluorescence (IF) staining for BrdU incorporation, p16^INK4A^ and p21^CIP1^ following 5 days of 4OHT treatment. n=5 biological replicates from 2 independent experiments. **g-j.** Relative mRNA expression for proinflammatory SASP components *(IL1A, IL1B, IL8 and CCL20)* assessed by RT-qPCR-following 4 days of 4OHT treatment with the indicated siRNAs (siControl, siS6K1_2, siS6K1_4, siS6K2_4 and siS6K2_5 n=4 for *IL1A, IL1B, IL8* and n=3 for CCL20; siS6K1/2 and siC/EBPβ n=3 for *IL1A, IL1B, IL8* and CCL20) in IMR90 ER: RAS cells. mRNA expression was normalized to *Rps14* housekeeping gene. *n* denotes independent experiments. **k.** Immunoblot analysis of S6K1, S6K2, IL-1β, IL-8 and GAPDH following 7 days of 4OHT treatment with the indicated siRNAs in IMR90 ER: RAS cells. IL-8 and GAPDH (loading control) were run on the same blot. S6K1, S6K2 and IL-1β were run on separate blots. Data are expressed as mean ± SEM. Statistical significance was calculated using one-way analysis of variance with Dunnett’s multiple comparison test (d-k, o-t) (* P < 0.05, ** P < 0.01, *** P < 0.001, **** P < 0.0001). n.s: non-significant.

Knocking down *S6K1, S6K2* or both kinases did not prevent senescence growth arrest as evaluated by measuring BrdU incorporation (Fig. 5d) or by quantification of the expression of key mediators of senescence growth arrest such as the CDKI p16^INK4a^ and p21^CIP1^ (Fig. 5e-f). Interestingly, the transcriptional induction of SASP components (*IL1A, IL1B, IL8*, *CCL20* and IL6, Fig. 5g-j and Extended Data Fig. 6c) was reduced upon knockdown of *S6K1* and/or *S6K2* with greatest effect observed with combined depletion. Immunoblot analysis confirmed that knocking down *S6K1* and *S6K2* resulted in reduced SASP expression (Fig. 5k).

Ribosomal protein S6 is the best-known target for S6K1 and S6K2 that phosphorylate it on serine residues 240 and 244 ^7^. Knocking down of both *S6K1* and *S6K2* resulted in a significant decrease of S6^S240/S244^ phosphorylation as assessed by IF (Extended Data Fig. 6d-e). Interestingly, individual knockdown of S6K1 and S6K2 had no or minimal effect on S6^S240/S244^ phosphorylation (Extended Data Fig. 6d-e), suggesting that S6^S240/S244^ phosphorylation might not explain the inhibitory effects on the SASP observed with S6K1 and/or S6K2 knockdown.

To further investigate this point, we took advantage of the S6K inhibitor LY2584702 (Extended data Fig. 7a). Treatment with LY2584702 resulted in a dose dependent inhibition of pS6^S240/S244^ but not 4EBP1 phosphorylation (Extended Data Fig. 7b-e). Treating IMR90 ER:RAS cells with LY2584702 did not rescue the senescence growth arrest as observed in colony formation assays or measuring BrdU incorporation, although it resulted in a trend of fewer cells positive for SA-β-Gal, similar to what was previously observed using the mTOR inhibitor Torin1 ^25^ (Extended Data Fig. 7f-h). Treatment with the S6K inhibitor caused a slight reduction of expression of SASP components such as IL1A or IL1B, not comparable to that observed with mTOR inhibition or depletion of *S6K1* and/or *S6K2* (Extended Data Fig. 7i-j). The above results show that knockdown of S6K in human cells decreased production of proinflammatory SASP components without preventing senescence.

### Transcriptional analysis shows that S6K1 regulates inflammatory pathways

To further analyse the relationship between S6K1 and the inflammatory SASP, we carried out RNA-Seq analysis of MEFs undergoing serial passage or *Ras*-induced senescence (Fig. 6a). GSEAs showed that signatures related to inflammatory cytokines and/or interferon were upregulated in late passage WT MEFs (Fig. 6b) and upon *Ras* expression in these cells (Fig. 6c). Interestingly, these signatures were downregulated when comparing *S6K1* KO and WT MEFs (Fig. 6b-c). A similar downregulation of inflammatory and interferon-related signatures was also observed when comparing the transcriptional profile of late passage and Ras expressing *S6K1/S6K2* DKO and WT MEFs (Extended Data Fig. 8a-c). In agreement with these observations, multiple pro-inflammatory SASP components were downregulated in *S6K1* KO (Fig. 6d) and *S6K1/S6K2* DKO (Extended Data Fig. 8d) MEFs undergoing Ras-induced senescence when compared with their WT counterparts. To further explore how *S6K1* loss could result in decreased expression of inflammatory mediators *in vivo*, we conducted an Ingenuity pathway analysis (IPA) of the transcriptional profile of livers of old WT and *S6K1* KO mice and compared them with those of MEFs of the same genotypes undergoing Ras-induced senescence (Fig. 6e). Biological function analysis, confirmed that *S6K1* deletion in mice was associated with a downregulated inflammatory response and other associated functions such as reduced chemotaxis or leukocyte migration (Fig. 6f). Search for upstream regulators identified several components of the IFN pathway, including IFNg, Ifnar and IRF3. CGAS and STING were also identified as upstream regulators (Fig. 6f). Interestingly, S6K1 (and S6K2) has been previously proposed to interact with STING to activate IRF3, a key mediator of inflammatory responses, in a kinase-independent manner ^27^. Therefore, the above results suggest that S6K1-mediated modulation of the STING/IRF3 axis in a kinase-independent manner might explain the decreased expression of pro-inflammatory factors in cells and mice of *S6K1* KO genotype observed during ageing and senescence.

**Figure 6.**
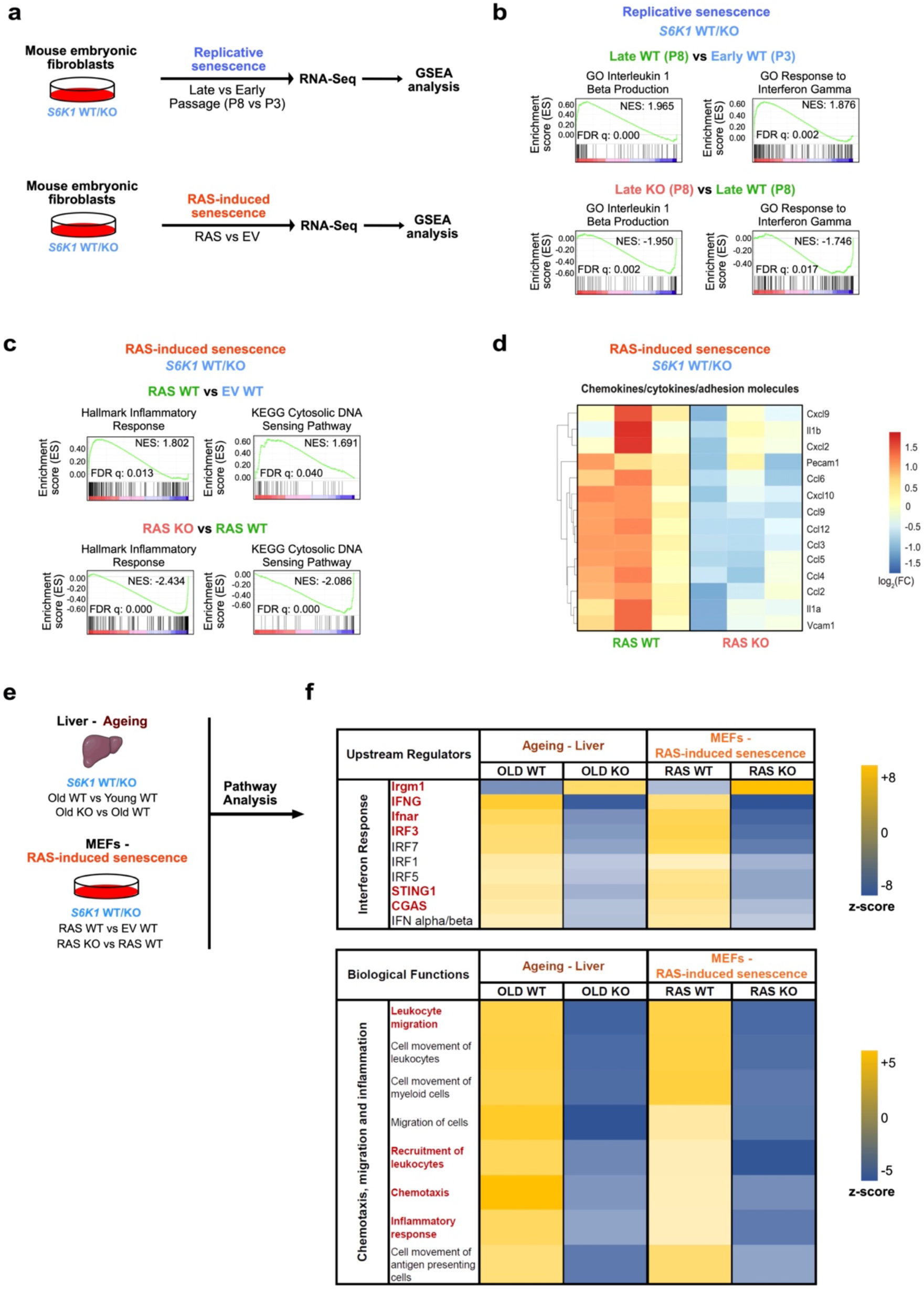
Transcriptional analysis shows that S6K1 regulates inflammatory pathways. **a.** Experimental scheme. Mouse embryonic fibroblasts (MEFs) from *S6K1* WT/KO embryos were assessed for replicative senescence or RAS-induced senescence. Samples underwent subsequent RNA-sequencing and gene-set enrichment analysis (GSEA). **b.** GSEA of early *S6K1* WT (passage 3), late *S6K1* WT (passage 8) and late *S6K1* KO (passage 8) MEFs. **c.** GSEA of *S6K1* WT MEFs expressing an empty vector (EV), *S6K1* WT MEFs expressing RAS^G12V^ or *S6K1* KO MEFs expressing RAS^G12V^. **d.** Heatmap illustrating the gene expression pattern of key proinflammatory SASP factors involved in RAS-induced senescence. **Left:** comparison of *S6K1* WT MEFs expressing RAS^G12V^ (n=3) with *S6K1* WT MEFs expressing EV (n=3). **Right:** comparison of *S6K1* KO MEFs expressing RAS^G12V^ (n=3) with *S6K1* WT MEFs expressing RAS^G12V^ (n=3). **e.** Schematic of combined pathway analysis of the aging cohort and in MEFs undergoing RAS-induced senescence of the indicated comparisons to identify common upstream regulators and biological functions. **f. Top:** assessment of common upstream regulators of the SASP in *S6K1* KO mice in the aging liver and in *S6K1* KO MEFs undergoing RAS-induced senescence. **Bottom:** assessment of biological functions that are commonly regulated in *S6K1* KO mice in the aging liver and in *S6K1* KO MEFs undergoing RAS-induced senescence. NES: normalized enrichment score. FDR: false discovery rate. WT: wild type. KO: knockout.

### S6K1 regulates senescence surveillance

To confirm that S6K1 regulates inflammatory responses, affecting leukocyte chemotaxis/migration through S6K1, we took advantage of a well-described mouse model of OIS and senescence surveillance. (Fig. 7a). In this model, hydrodynamic tail vein injection (HDTVi) of a transposon vector expressing oncogenic Nras (Nras^G12V^), induces OIS and triggers a SASP-dependent immune surveillance response that causes the clearance of preneoplastic hepatocytes ^40^.

We analysed the Nras+ cells present 4 days and 7 days after HDTVi in the livers of *S6K1* WT and *S6K1* KO mice. At day 4, before immune-mediated clearance of senescent cells occurs, the numbers of Nras+ cells were similar in mice of both genotypes. The number of Nras+ cells decreased in mice of both genotypes by day 7, but there was significantly higher number of Nras+ cells in *S6K1* KO mice when compared with their WT counterparts at day 7 (Fig. 7b-c). These differences were likely due to an impaired immune-mediated elimination of senescent cells in *S6K1* KO mice, as myeloid (major histocompatibility complex II, MHC-II+, CD68+) and lymphoid (CD3+) infiltrates observed in the liver of WT mice at day 7 were significantly lower in *S6K1* KO mice (Fig. 7d-i).

**Figure 7.**
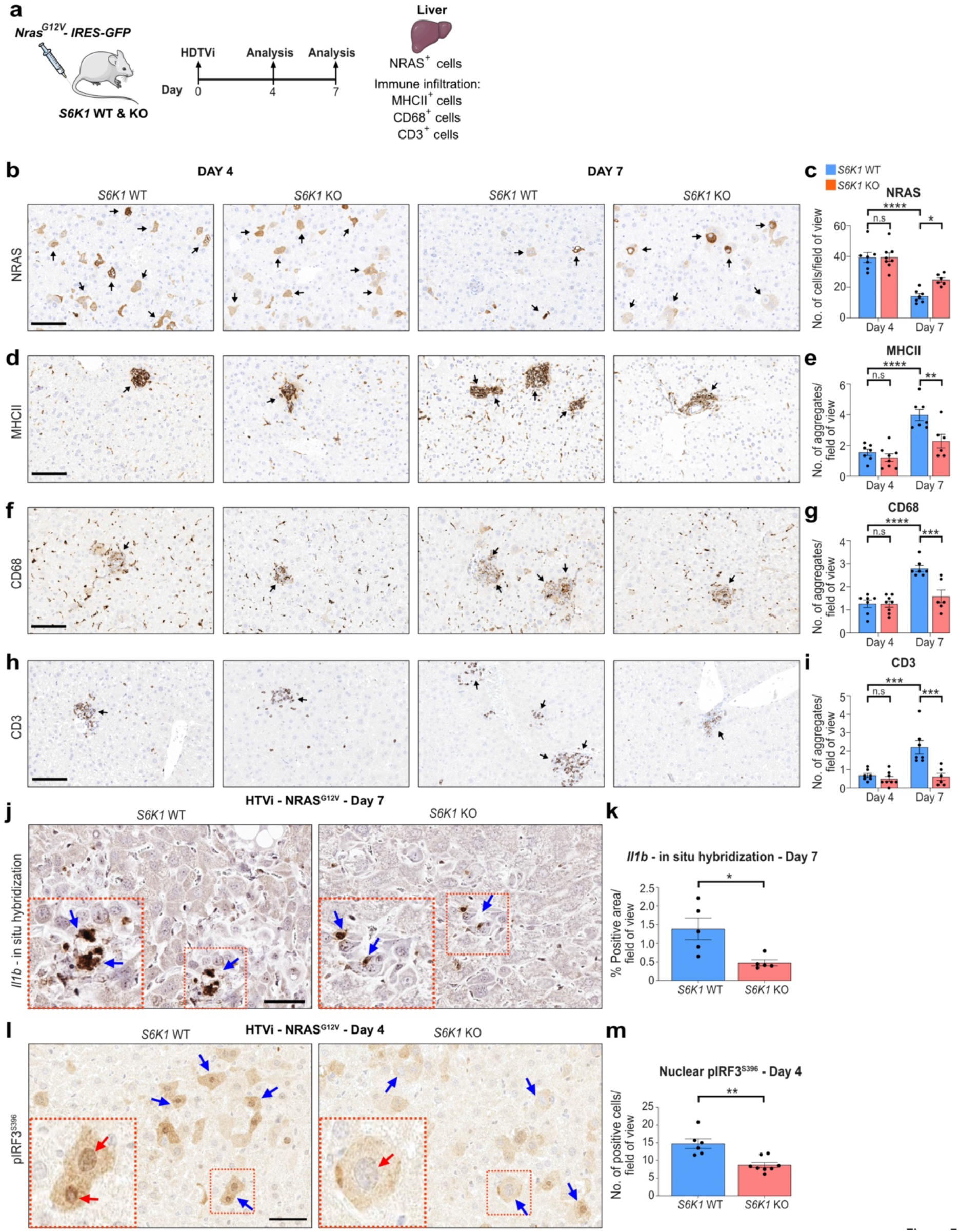
S6K1 regulates senescence surveillance. **a.** Experimental scheme. Hydrodynamic tail vein injection (HDTVi)-based co-delivery of an *Nras^G12V^* transposon construct and a transposase expressing vector into mouse livers (day 0). Mice were sacrificed 4 or 7 days following HDTVi to assess senescence surveillance. **b-i.** Immunohistochemistry staining for NRAS, MHCII, CD68 and CD3 **(left)** and quantification **(right)** of livers from Day 4 *S6K1* WT (n=7), KO (n=8) mice and in Day 7 *S6K1* WT (n=7), KO (n=6) mice. **j, k.** *In situ* hybridization for *Il1b* mRNA **(left)** and quantification **(right)** of livers from Day 7 *S6K1* WT (n=5) and KO (n=5) mice. **l, m.** Immunohistochemistry staining for pIRF3^S396^ **(left)** and quantification **(right)** of livers in Day 4 *S6K1* WT (n=6) and KO (n=8) mice. Data are expressed as mean ± SEM. Statistical significance was calculated using two-way analysis of variance with Tukey’s multiple comparison test (c, e, g, i) or by Student’s t-test (k, m). (* P < 0.05, ** P < 0.01, *** P < 0.001, **** P < 0.0001). *n* denotes individual mice. n.s: non-significant.

To investigate whether these differences in infiltration were due to decreased expression of immunomodulatory SASP components in the senescent hepatocytes, we performed *in situ* mRNA hybridization to detect transcripts for *Il1b* at day 7 (Fig. 7j). *Il1b* is one of the cytokines more robustly regulated by S6K1 and plays a critical role in activating immune surveillance of senescent cells ^41^. Consistent with our previous observations, there was less *Il1b+* cells upon oncogenic Ras expression in *S6K1* KO mice (Fig. 7k). To understand if the reduced expression of immunomodulatory SASP components could be due to S6K1-mediated modulation of the STING/IRF3 axis ^27^, we examined the expression of nuclear IRF3 phosphorylated on Ser396 (pIRF^S396^), which indicates activation of the STING/IRF3 axis. While we could observe many hepatocytes with nuclear pIRF^S396^ in WT mice, we often observed only cytoplasmic staining in *S6K1* KO mice (Fig 7l). The significance of this observation was confirmed upon quantification of nuclear pIRF3 (Fig 7m). The above results suggest that impaired activation of the STING/IRF3 axis in *S6K1* KO mice might explain, at least in part, the reduced inflammation observed in these mice.

In conclusion, we have observed that S6K1 regulates age-related inflammation (inflammaging) and senescence surveillance through modulation of key proinflammatory chemokines/cytokines. Given that chronic inflammation is a driver of many age-related pathologies, these results may explain why *S6K1* KO mice are long-lived and show improved healthspan.

## DISCUSSION

Inhibition of mTOR signalling, including the key effector S6K1 extends lifespan in an evolutionarily conserved manner and increases healthspan in mice ^9^. Underlying mechanisms include beneficial long-term effects on glucose homeostasis, adipose tissue biology and effects upon key hormone and energy sensing signalling pathways ^9, 12, 13, 17^. In the current study, we demonstrate that loss of S6K signalling in the liver has marked anti-inflammatory effects and attenuates various age-related hepatic liver pathologies. This blockade of S6K signalling appears to act at least in part via changes in age-related inflammation (inflammaging) rather than altering senescence *per se* with concomitant beneficial effects on associated phenotypes such as age-related fibrosis. Furthermore, our studies aimed at further unravelling the cellular and molecular mechanisms at work demonstrate that deletion of S6Ks intrinsically impairs the production of proinflammatory cytokines in both mouse and human cells, has a broad effect on the senescence associated inflammatory profile and immune cell recruitment. At a molecular level, our data suggests that at least in part the effects seen in the liver are related to a non-kinase dependent role of S6Ks in regulating STING/IRF3 ^27^, a key modulatory axis of immune activation.

Increased accumulation of senescent cells is a recognised feature of ageing in the mouse liver although its precise origin and its pathophysiological impact remains to be determined ^31, 35^. Work by others and us has shown that inhibition of mTOR signalling leads to a specific inhibition of the proinflammatory SASP while not affecting senescence growth arrest, an effect in part mediated by 4EBP ^25, 26^. However, these studies had not specifically investigated the role of S6Ks in these findings even though loss of S6K1 has beneficial effects on ageing and related liver phenotypes ^9, 32^. Our current studies demonstrate *in vivo* and in multiple cellular models, that depletion of S6K1 and 2 (singly or in combination) does not alter the accumulation of senescent cells or the senescence response *per se* but rather affects their pro-inflammatory properties.

We show that loss of S6K signalling has marked effects upon the SASP with a profound reduction in a subset of inflammatory markers. In the liver of old S6K1 KO mice, several inflammatory cytokines showed reduced expression and this was associated with lowered immune cell infiltration. Ageing in the liver is associated with low grade inflammation, which may in part be related to calorie and macronutrient intake ^31, 35^. Accumulation of senescent cells in the liver has been reported to promote hepatic fat accumulation and steatosis features of liver ageing and ablation of these cells ameliorates this phenotype ^35^. Our findings suggest that the primary driver of the beneficial effects of the removal of senescent cells upon late life metabolic dysfunction in the liver stems from the abrogation of the proinflammatory profile engendered by these cells.

Systemic deletion/inhibition of S6K1 and/or S6K2 has beneficial effects upon lipid accumulation in the liver in states of overnutrition as well as having beneficial effects upon a range of age-related liver phenotypes ^9, 42, 43^. However, due to the cross talk between the liver and key metabolic tissues such as adipose tissue and skeletal muscle and the known effects of S6K1 deletion in these tissues, it is unclear whether beneficial effects of this manipulation stem from liver intrinsic effects. Further work will be needed to investigate this question but existing studies in mice with virally mediated loss of S6K suggest that liver intrinsic effects of S6K loss have beneficial systemic metabolic effects ^32^ although the role of senescence and the impact upon longevity were not studied.

Our previous studies have demonstrated that global deletion of S6K1 results in beneficial effects on longevity and healthspan ^9^. S6K1 belongs to a family of S6Ks which also includes S6K2 which remains an understudied signalling component ^44^. S6K1 and S6K2 share common substrates but also have specific ones ^7^. Combined global loss of S6K1 and S6K2 protects mice from the negative effects of high fat diet^42^ but the individual and/or combined contribution of each kinase to processes such as senescence and the SASP were unknown.

Our in vitro studies suggest that while genetic knockdown of S6K1 resulted in alterations in a range of parameters such as the reduction in inflammatory markers, deletion of S6K2 alone had limited effects. An exception was in OIS in human cells where KD of S6K2 had significant effects upon inflammatory markers suggesting that S6K2 could have specific roles in this process or play a more important role in human cells compared to mouse cells. In general, combined deletion of both kinases had the most marked effects on inflammatory mediators in both mouse and human cells and across the range of cell models of senescence that we studied.

Surprisingly LY2584702, a pharmacological inhibitor of S6K ^45^, which was effective in blocking S6K action as judged by the phosphorylation of S6, had only a minimal effect on OIS-induced inflammation. This hints at the concept that non-kinase-dependent effects of S6K1 might mediate, at least in part, the anti-inflammatory effects of abrogating this pathway. Indeed, it has previously been shown that a tripartite interaction between S6K1, STING and TBK1 mediates the activation of the transcription factor IRF3, a key component in early innate immune responses ^27^. Importantly, this was independent of S6K1 kinase function ^27^. While LY2584702 has been shown capable to mitigate diet-induced hepatosteatosis ^43^, our results suggest that a PROTAC derivative able to degrade S6K could be a more effective alternative. IRF3 is also involved in DNA damage associated cell senescence ^46^, induction of the SASP, and potentially plays a role in premature ageing at least in cellular models ^47^. Based on these observations, we explored a potential role for the IRF3 and found impaired activation of this molecule in OIS in the livers of S6K1 KO mice. This finding might underlie in part the reduced inflammation seen in these mice. S6Ks have multiple substrates, however, and future work will explore the role of these in the regulation of the SASP.

In summary our findings show that loss of S6K signalling both in ageing models *in vivo* and *in vitro* attenuates senescence associated inflammatory processes and may play an important role in the beneficial effects of attenuation of mTOR signalling on detrimental phenotypes particularly in the ageing liver.

## METHODS

### Mice experiments

All mice were kept under specific pathogen-free barrier conditions within individually ventilated cages on a 12-hour light/dark cycle between 21-23°C. Mice were given *ad libitum* access to food and water. Mice were fed chow (RM3 expanded; Special Diets Services) diet. *S6K1* KO or *S6K2* KO mice on a C57BL/6 inbred strain were described previously ^9, 48^. The experiments were performed in accordance to the UK Animals (Scientific Procedures) Act 1986 and amended regulations (2012) and approved by the Imperial Collegés animal welfare and ethical review body under either 70/8700 or 70/09080.

Ageing experiment. Female *S6K1* wild type (WT) and knockout (KO) mice were generated from heterozygous breeding pairs or trios and aged for either 90 days (young) or 600 days (old) prior to being sacrificed.

Hydrodynamic tail vein injection (HDTVi) experiment. Male 8-10-week-old *S6K1 WT* and *KO* mice were used. HDTVi was carried out to deliver transposon-based vectors as previously described ^40^. All vectors were prepared with the GenElute HP Endotoxin-Free Plasmid Maxiprep Kit (Sigma). On day 0, 25 µg of the vector expressing *Nras^G12V^* and 5 µg of the SB-13 Sleeping Beauty transposase expression vector were diluted in sterile PBS to a total volume of 2 mL (∼10% body weight) prior to HDTVi within 10 seconds. Livers were collected 4 and 7 days following the HDTVi.

### Mouse embryonic fibroblast (MEFs) generation

MEFs were prepared as previously described ^49^. Briefly, MEFs were prepared from 13.5-day embryos of mice bred in heterozygosity for S6K1 or S6K2. Importantly, both WT and KO MEFs of the indicated genotypes were prepared from the same mother to ensure littermates were utilized. MEFs generated from at least 3 independent mothers were used for experiments. *S6K1/S6K2* DKO MEFs were prepared by breeding mice that were both heterozygous for *S6K1 (S6K1^+/-^)* and knockout for *S6K2 (S6K2^-/-^)* mice together (*i.e., S6K1^+/-^; S6K2^-/-^ x S6K1^+/-^; S6K2^-/-^*). WT MEFs were used as controls for *S6K1/S6K2* DKO experiment. WT MEFs were prepared from embryos of WT mothers that were obtained at earlier stages of breeding for generation of double deletion.

Preparation of MEFs were performed by first removal of the embryo from the uterus and yolk sac followed by removal of the head and viscera. The remaining tissue was minced and triturated in trypsin-EDTA (0.05%, Gibco, USA) using a scalpel and gentle pipetting and incubation at 37°C and 5% CO2 for 15 minutes with periodic (every 5 minutes) resuspension. A single-cell suspension was then obtained by passing cells through a 100 μm sterile nylon cell strainer (Falcon, USA). Cells were cultured for 3-5 days until confluence was reached and were then frozen in complete DMEM (see below) with 10% dimethyl sulfoxide (DMSO; Sigma).

### Chemical compounds and drug treatments

4-hydroxytamoxifen (4OHT – 125 nM; Sigma-Aldrich, USA) was dissolved in DMSO. LY2584702 (2 μM, Key Organics, UK) and Torin1 (25 nM, Tocris, UK) were also dissolved in DMSO. Cells were treated with the indicated drugs the day after seeding. 4OHT was replenished every four days and LY2584702 and Torin1 were replenished every two days.

### Plasmids and vectors

pLNC-ER: RAS retroviral vector has been previously described ^50^. pBABEpuro empty vector (EV) or MSCV-neo vector expressing constitutively active RAS (HRAS^G12V^) has been previously described ^51^. EcoHelper (pCL-Eco, Addgene).

### Cell culture and retroviral transduction

MEFs were maintained in DMEM supplemented with EmbryoMax® FBS (Millipore, USA) and 1x Antibiotic-Antimycotic at 37°C and 5% CO2. Human IMR90 fibroblasts, human embryonic kidney 293 transformed (HEK293T) cells were obtained from the American Type Culture Collection (ATCC). For proliferation and maintenance of IMR90 and HEK293T lines, cells were grown in Dulbecco’s Modified Eagle Medium (DMEM) supplemented with 10% foetal bovine serum (FBS, Sigma-Aldrich, USA), 1x Antibiotic-Antimycotic (Gibco, USA) and kept incubated at 37°C and 5% CO2. Cells were cultured in the indicated medium during experiments unless otherwise stated. The Guava Viacount reagent (Millipore, USA) and the Guava Cytometer (Millipore, USA) were used to assess cell number and viability Cells were routinely assessed for mycoplasma.

Retroviral transduction was performed as previously described ^51, 52^. HEK293T cells with or without GagPol expression were used for packaging of retrovirus. For generation of IMR90 ER: RAS^G12V^ fibroblasts, transfection was performed in 10 cm dishes using HEK293T cells with GagPol expression, pLNC-ER: RAS vector and packaging vectors using 1mg/mL of linear 25 kDa polyethylenimine (PEI, Polysciences). 24 hours following transfection, medium was replaced with fresh 6 mL (to concentrate the virus) of complete DMEM and transfection efficiency was monitored by expression of mCherry using an Olympus CKX41 inverted light microscope. Human IMR90 fibroblasts were seeded at a density of 10^6^ per 10 cm dish on the same day. 48 hours following transfection, the viral supernatant was filtered (0.45 μm), supplemented with 4 μL of 8 mg/mL of polybrene and added to IMR90 fibroblasts for 3 hours. 2 additional rounds of transduction were carried out before replacing with fresh complete DMEM. 48-72 hours following transduction, cells were split and selected with Geneticin® (400 μg/mL).

For generation of MEFs expressing either empty vector (EV) or constitutively expressing HRAS^G12V^, transfection was performed as above with a few differences. Transfection was carried out using HEK293T cells in 10 cm dishes with viral vector, EcoHelper (pCL-Eco, Addgene) and packaging vectors using 1mg/mL 25 kDa PEI. 1.5×10^6^ MEFs were seeded for transduction. Transduction was carried out by pooling the empty vector or HRAS^G12V^ supernatant together and adding an equal amount of virus titre (6 mL) to each 10 cm MEF dish. Only a single 8-hour round of transduction was carried out. 48-72 hours following transduction, MEFs were selected with 3 μg/mL of puromycin and transduction efficiency for mCherry was assessed by flow cytometry using Guava EasyCyte™ (Millipore, USA). A transduction efficiency of 95% or greater was achieved.

### Reverse transfection of small interfering RNAs (siRNAs)

Lyophilized siRNAs targeting *S6K1* or *S6K2* were obtained from Qiagen in Flexitubes® with a preference for verified siRNA sequences. siRNAs were first reconstituted in RNase-free water to a concentration of 1 μM and aliquoted. Please refer to Supplementary Table 1 for siRNA sequences.

For RNA analysis, 1.2×10^5^ (for growing cells that will not be given 4OHT) or 2.4×10^5^ IMR-90 ER: RAS fibroblasts in suspension were reverse-transfected with the indicated siRNAs in a 6 cm dish to a final volume of 4 mL in DMEM with 10% FBS but without antibiotic. The transfection mix consisted of 4 μL of DharmaFECT1™ (GE Healthcare, UK), 144 μL of 1 μM siRNA (35 nM final concentration) and 700 μL plain DMEM. Each transfection mix was briefly vortexed and incubated at RT for 30 minutes prior to cell seeding. The transfection medium was replaced with fresh complete medium with or without 4OHT 16 hours later once cells had adhered. Allstars scrambled siRNA served as a negative control.

For high-content immunofluorescence analysis in 96-well format, 1.75×10^3^ (for day 5 analysis) or 1.25-1.5×10^3^ (for day 8 analysis) IMR90 ER: RAS fibroblasts in suspension were reverse transfected with the indicated siRNAs in a final volume of 100 μL in DMEM with 10% FBS but without antibiotic. The transfection mix consisted of 0.1 μL of DharmaFECT1™ (GE Healthcare, UK), 3.6 μL of 1 μM siRNA (35 nM final concentration) and 17.5 μL plain DMEM. Each transfection mix was briefly vortexed and incubated at RT for 30 minutes prior to cell seeding. The transfection medium was replaced with fresh complete medium with or without 4OHT 16 hours later once cells had adhered. Allstars scrambled siRNA served as a negative control.

### MEFs serial passage

Passage 1 MEFs of the indicated genotypes were seeded (2×10^6^ cells) in 10 cm dishes. Cell counts were performed using the Guava Cytometer (Millipore, USA) every 4 days (1 passage) until WT cells reached replicative exhaustion. Experiments were performed in 21% O2 and medium was replaced every 2 days. Cumulative population doublings per passage was calculated as log2 (number of cells at time of subculture/ number of cells plated) and plotted against the total time in culture (passage 2 until 8). RNA was extracted from cells at passages 3 (young) and 8 (old). Cells were also seeded to assess 5-bromo-2’-deoxyuridine (BrdU) incorporation and senescence-associated beta galactosidase (SA-β-Gal) activity at passages 2 and 8.

### Crystal violet staining

Cells were seeded (1.5-2×10^4^) in 6-well dishes. Cells were cultured until control (growing) cells reached confluence (usually 13 days). Every 2-4 days the medium was replenished with the appropriate drug treatments. Cells were fixed with 0.5% glutaraldehyde (v/v) (Sigm-Aldrich, USA) and stained with 0.2% crystal violet (w/v) (Sigma-Aldrich, USA).

### SA-β-Gal staining

SA-β-Gal activity was assessed as previously described ^53^. For cytochemistry assays, passage 2 and 8 MEFs were seeded (8×10^4^) in 6-well dishes and fixed the following day with 0.5% glutaraldehyde (v/v) (Sigm-Aldrich, USA). Cells were washed twice in 1 mM magnesium chloride (MgCl2) in PBS (pH 5.5) and incubated in X-gal staining solution (1mg/mL X-Gal, Thermo Scientific, 5 mM K3[Fe(CN)6] and 5 mM K4[Fe(CN)6*3H2O]) for 8 hours at 37°C. Bright-field images were acquired using the Olympus CKX41 inverted light microscope and the Olympus DP20 digital camera. The percentage of SA-β-Gal positive (blue staining) cells was estimated by counting at least 150 cells per well.

For fluorescence assays, IMR90 ER: RAS cells (1.5-2.5×10^3^) were seeded in 96-well plates in triplicate and treated with the indicated drugs on the following day. 8 days later, cells were incubated with fresh DMEM containing DDAO galactoside (9*H*-(1,3-dichloro-9,9-dimethylacridin-2-one-7-yl) β-D-Galactopyranoside) (Molecular Probes™) for 2 hours. Cells were then fixed in 4% formaldehyde solution (v/v) (Sigma-Aldrich, USA), washed and stained with 4’, 6-diamidino-2-phenylindole (1 μg/mL DAPI) for 15 minutes. Fluorescence images were acquired and analysed by high-content analysis microscopy using the InCell Analyzer 2000 (GE Healthcare, UK) and the InCell Investigator software 2.7.3. The percentage of SA-β-Gal positive (blue staining) cells was estimated by counting at least 1000 cells per well.

### Assessing BrdU incorporation

BrdU incorporation was assessed as previously described ^53^. Briefly, BrdU incorporation was carried out by high-content microscopy and analysis.

IMR90 ER: RAS fibroblasts were seeded (1-3×10^3^) in 96-well plates in triplicate, allowed to adhere over-night and treated with or without 4OHT (± any additional treatments) on the following day. Cells were then pulsed overnight (17 hours) in BrdU (50 μM) and then fixed in 4% formaldehyde solution (v/v). Cells were permeabilised with 0.2% Triton X-100 (v/v) (Sigma-Aldrich, USA) for 15 minutes and blocked (0.5% BSA (w/v) and 0.2% fish skin gelatin (v/v) in PBS) for 1 hour at room temperature (RT) with gentle shaking. Fixed cells were incubated with mouse anti-BrdU primary antibody, DNAse (0.5 U/μL, Sigma-Aldrich, USA) in the presence of 1 mM MgCl2 in blocking solution and incubated for 30 minutes at RT. Cells were washed prior to incubation with goat anti-mouse Alexa Fluor 594® secondary antibody in blocking solution for 30 minutes. Cells were washed prior to incubation with 1 μg/mL DAPI. Plated were then analysed by high-content microscopy (described below).

Passage 2 and 8 *S6K1/S6K2* WT/DKO MEFs for the cumulative population doubling experiment were seeded (8×10^3^) in 96-well plates in triplicate and allowed to adhere over-night. Whereas passage 2 MEFs from S6K1 WT/KO, S6K2 WT/KO and/or S6K1/S6K2 WT/DKO MEFs for the time course were seeded at a density of 3×10^3^ in 96-well plates in triplicate and allowed to adhere-overnight. Cells were pulsed for 8 hours with BrdU (50 μM) and processed as described above.

### Immunofluorescence (IF) and high-content analysis (HCA) microscopy

IF and HCA were performed as previously described ^25, 53^. A list of antibodies and dilutions used can be found in Supplementary Table 2. Cells were seeded in 96-well plates (Nunc, Thermo Fisher Scientific, USA), allowed to adhere overnight, treated with the indicated drugs on the following day and fixed on the desired day of analysis. Medium was replaced every 2-4 days. Cells were fixed in 4% formaldehyde solution (v/v), permeabilised with 0.2-0.3% Triton X-100 (v/v) and blocked with 0.5% BSA (w/v) and 0.2% fish skin gelatin (v/v) in PBS. For SASP and RPS6 staining, cells were blocked in a solution containing 5% donkey serum, 0.3% Triton X-100 and 0.1% bovine serum albumin (BSA). Cells were incubated with primary antibody for 1 to 1.5 hours hours at RT, washed followed by incubation with the appropriate secondary antibody (Alexa Fluor 488® and/or 594® - 1:750 dilution). Cells were then washed prior to incubation with 1 μg/mL DAPI.

Images were acquired using an automated-high-throughput microscope (InCell Analyzer 2000, GE Healthcare, UK) with a 20X objective. Image acquisition was set up so that at least 1000 cells/well from multiple fields were detected. The InCell Analyzer 2000 captured images in 4 different wavelengths (DAPI, FITC – 488, Texas Red – 594 and PE-Cy5 – DDAOG). Experiments were performed in either duplicate or triplicate wells. The InCell Investigator software 2.7.3 (GE Healthcare, UK) was used for image processing and quantification. Nuclei segmentation and cell identification was performed using DAPI staining. Nuclei were segmented using a top hat segmentation approach (a minimum area of 120 μm^2^). The cell area was defined using either a collar segmentation approach that placed a border of 2 μm around the DAPI staining or alternatively using cytoplasmic intensity by a multiscale top-hat approach. The cellular expression (nuclear or cytoplasmic) of the protein of interest was calculated by quantifying the mean intensity of pixels in the desired reference channel (FITC 488 or Texas Red 594). A histogram was generated that assigned intensity values for all of the cells in a given sample. Then a threshold filter to define the number of positive and negative cells for the given protein or signal of interest was set up by assigning a nuclear or cytoplasmic intensity value to each cell that correlates to the specific expression. Alternatively, normalized intensity values for a given staining was calculated by measuring the difference between the raw intensity and intensity of the background (secondary antibody alone). The relative fold change to a specified condition (*e.g.,* cells treated with 4OHT) was then calculated using the normalised intensity values. Specificity of antibodies was validated with the use of robust controls (shRNAs/siRNAs, overexpression or drug inhibition).

### Immunohistochemistry (IHC)

Tissues were fixed in 4% paraformaldehyde solution (Santa Cruz Biotechnology, USA) overnight at 4°C, dehydrated and embedded in paraffin. Paraffin-embedded liver sections (2 μm or 7 µm for Sirius-Red staining) were processed for IHC staining using a BOND-MAX system (Leica, Germany) as previously described ^54^. Following deparaffinization and rehydration, antigen retrieval was carried out with Bond^TM^ citrate solution (AR9961, Leica), Bond^TM^ EDTA solution (AR9640, Leica) or Bond^TM^ proteolytic enzyme kit (AR9551, Leica). Sections were then incubated with the indicated antibodies in Bond^TM^ primary antibody diluent (AR9352, Leica Biosystems). This was followed by incubation with secondary antibodies (Leica) and staining using the Bond Polymer Refine Detection Kit (DS9800, Leica). Whole slides were then scanned using an Aperio AT2 slide scanner (Leica, Germany) at 20x objective. Whole slides were annotated and analyzed using Aperio ImageScope (ver. 12.4.0.5043, Leica) and Fiji (ImageJ ver. 1.52e, National Institutes of Health USA).

Antibodies that were used are anti-Ki67, rabbit, 1:200 (Thermo Scientific, RM-9106-S1); anti-CHOP, rabbit, 1:100 (Cell Signaling, #5544); anti-BiP, rabbit, 1:200 (Cell Signaling, #3177); anti-MHCIl, rat, 1:500 (clone M5/114.15.2, Novus Biologicals, NBP1-43312); anti-CD68, rabbit, 1:200 (abcam, 125212); anti-F4/80, rat, 1:250 (Linaris, T2006); anti-CD3, rabbit, 1:500 (clone SP7, Invitrogen, MA1-90582), anti-B220, rat, 1:3000 (clone RA3-6B2 – BD Biosciences, 553084), anti-CD4, rat, 1:1000 (eBioscience, 14-9766); anti-CD42b, rabbit, 1:200 (Abcam, clone SP219, ab183345). anti-NRAS, mouse, 1:50 (Santa Cruz - sc-31), anti-pIRF3^S396^, rabbit, 1:300 (Bioss, BS-3195R).

### RNA *in situ* hybridization

RNA *in situ* hybridization in 6 μm liver sections was performed using the RNAscope® 2.5 Assay (FFPE and 2.5 HD Brown Assay) from Advanced Cell Diagnostics (ACD), according to manufacturer’s protocol. Probes for Mm-*Il1b* (Cat No. 316891), the housekeeping gene Mm-*Ppib* (positive control, Cat No. 313911) and the bacterial gene *dapB* (negative control, Cat No. 310043) were purchased from ACD. Signal detection was carried out by DAB staining. Slides were counterstained with haematoxylin prior to mounting and then whole digital slides were acquired using the Aperio AT2 slide scanner (Leica, Germany) at 40x objective.

### Peripheral complete blood count

Whole blood was collected from the tail vein using citrate as an anticoagulant and diluted using saline to a volume of at least 200 μl. Complete blood counts were then obtained using the Sysmex XE2100 automated cell counter (Sysmex Corporation, Japan).

### Immunoblotting

Snap frozen liver tissues (30 mg) were lysed and homogenised in RIPA lysis and extraction buffer (50 mM Tris pH8, 150 mM NaCl, 1% Triton-X 100, 0.5% Na-Doc, 0.1% SDS and 1 mM EDTA) containing cOmplete^TM^, Mini, EDTA-free protease and phosphatase inhibitors (Roche, Switzerland) using the IKA® T10 basic ULTRA-TURRAX homogeniser (IKA, Germany). Lysates were incubated on ice for 10 minutes for lysis prior to being spun at *16,000 g* (4°C) for 10 minutes. Supernatant (protein extracts) was collected. Cells were washed twice in ice-cold PBS prior to lysis in RIPA buffer.

Protein concentration was measured using the BioRad *DC*® Protein Assay (BioRad, UK) according to manufacturer’s protocol and samples were denatured in Laemmli sample buffer. Samples were loaded into a 4-15% Mini-PROTEAN TGX Precast Gel (BioRad, UK). Immunoblotting was carried out as previously described ^55^. Please refer to Supplementary Table 2 for the list of antibodies and dilutions used.

### Gene expression analysis

Total RNA was extracted from liver tissues or cells using the TRIzol® reagent (Ambion) and the RNeasy Mini Kit (Qiagen, Germany) as previously described ^53^. cDNA synthesis was carried out using SuperScript® II reverse transcriptase (Invitrogen, USA) kit with dTNPs and random hexamers according to manufacturer’s protocol. PCR reactions were carried out in a CFX96^TM^ Real-Time PCR system (Bio-Rad, USA) using Power SYBR Green Master Mix (Applied Biosystems, USA). Gene expression was normalized to ribosomal protein S14 (RPS14) housekeeping genes unless stated otherwise and analysed by the comparative Ct method. Please refer to Supplementary Table 3 for the list of primers used.

### RNA-sequencing and analysis

RNA purity and integrity were first determined with an Agilent 2100 Bioanalyser and the RNA 6000 Nano Kit high, and an RNA integrity score (RIN) above 7 was used for quality assurance.

The cDNA library was then prepared from 500 ng of total RNA using the TruSeq stranded mRNA library kit (Illumina, USA) according to the manufactureŕs protocol. The quality of the library was assayed with an Agilent 2100 Bioanalyser with a HS DNA chip and the DNA concentration was measured with a Qubit (Life Technologies, USA). The cDNA library was sequenced with a HiSeq 2500 (Illumina, USA) with single end, 50-base pair reads for a minimum of 40 million reads per sample.

The data were processed using standard procedures. In brief, demultiplexing was carried out with CASAVA software (ver. 1.8.4) and raw reads were mapped to the mm9 genome using TopHat aligner. The “HTseq counts module” was used to obtain gene-based counts. The “DESeq2” Bioconductor package was used for differential expression analysis. Sample-to-sample distances were calculated using the *“dist”* functions in “DESeq2” was employed to calculate. Gene set enrichment analysis (GSEA) was carried out using the Broad Institute GSEA application based on Wald statistics obtained from the “DESeq2” comparisons. Genes with very few read events were excluded. Distances were visualised by *‘pheatmap’* function available in ‘pheatmap R package’.

The Broad institute GSEA application was used to carry out Gene set enrichment analysis (GSEA). The genes were ranked based on “Wald statistics” available from ‘DESeq2’ comparisons. Genes with very few read counts were excluded from analysis. QIAGEN Ingenuity Pathway Analysis (IPA) was used to identify common “upstream regulators” and “biological functions”.

### Statistical analysis

Data are expressed as mean ± standard error of mean (SEM) unless stated otherwise. Statistical analysis was performed in Graph Pad Prism 9 software. Statistical significance for comparison of two groups was calculated using unpaired Student’s t-test. Comparison of three or more groups with one variable was calculated using one-way analysis of variance (one-wat ANOVA) with Tukey’s or Dunnett’s multiple comparison test. Comparison of four groups with two variables was done using two-way analysis of variance (two-way ANOVA) with Tukey’s multiple comparison test. A p-value of < 0.05 was considered statistically significant. * P <0.05, ** P< 0.01, *** P<0.001 and **** P<0.0001.

### Reporting summary

Further information on research design is available in the Nature Research Reporting Summary linked to this article.

## DATA AVAILABILITY

Source data will be included with the published manuscript.

## ACKNOWLEDGEMENTS

We are grateful to Tom Carroll, Virinder Reen, Athena Georgilis and other members of J. Gil’s laboratory for reagents, comments and other contributions to this project. We are thankful for Mirian Fernandez and Adnan Ali from M. Heikenwalder’s laboratory for their support. For the purpose of open access, the authors have applied a Creative Commons Attribution (CC BY) licence. Core support from MRC (MC_U120085810) funded this research in J. Gil’s laboratory. M. Heikenwalder was supported by an European Research Council (ERC) Consolidator grant (HepatoMetaboPath), SFB/TR 209 project ID 314905040, SFB 1479 (Project ID: 441891347), SFB1479, the Rainer Hoenig Stiftung, Research Foundation Flanders (FWO) under grant 30826052 (EOS Convention MODEL-IDI) and a seed funding from HI-TRON. Work was supported by the Medical Research Council grant MC-A654-5QB40 and a Wellcome Trust Grant 098565/Z/12/Z in D. J. Withers’ lab.

## AUTHOR CONTRIBUTIONS

S.G. conceived and designed the project; conceived, designed, performed and analysed experiments and aided with the manuscript writing. E.E.I designed and performed *in vivo* experiments. S.M.A.P designed and performed *in vivo* and *ex vivo* experiments. J.E.B.A. performed immunohistological staining and analysis. S.K carried out bioinformatics analyses. J.P. aided with in vivo experiments. S.B. aided with in vitro and in vivo experiments. D.H. performed immunohistological staining. G.D. carried out bioinformatics analyses. A.I.C. performed biochemical analyses. N.H performed *in vitro* experiments. S.V. designed and performed in vivo experiments. M.H. conceived and designed the project, secured funding and aided with manuscript writing. J.G. and D.J.W. conceived and designed the project, secured funding and wrote the manuscript, with all authors providing feedback.

