## Extended Data for "Ribosomal S6 kinase 1 regulates ‘inflammaging’ via the senescence secretome"

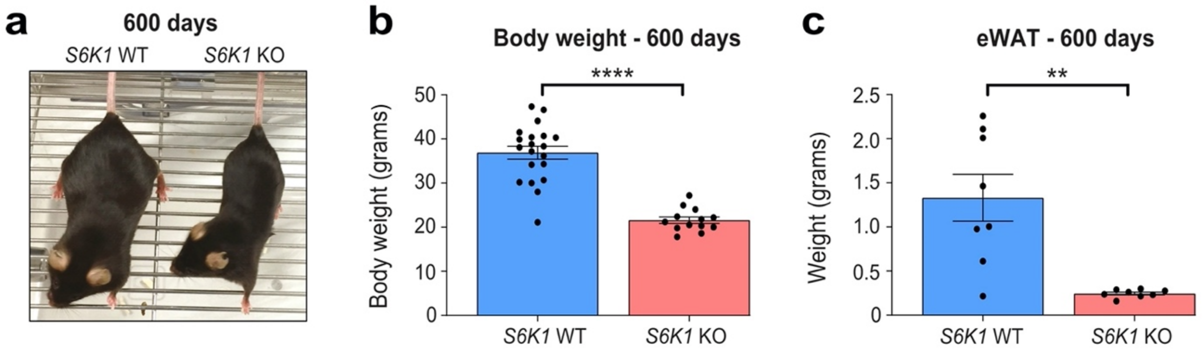


**Extended Data Figure 1. S6K1 deletion attenuated age-induced obesity.** *S6K1* wild type (WT) and knockout (KO) mice were aged for 600 days. a**.** Representative photograph of 600-day-old *S6K1* WT (left) and KO (right) mice. **b.** Body weight (grams) at 600 days of *S6K1* WT (left; n=20) and KO (right; n=13) mice. **c.** Epididymal white adipose tissue (eWAT) weight (grams) at 600 days of *S6K1* WT (left; n=8) and KO (right; n=8) mice. Data are expressed as mean ± SEM. Statistical significance was calculated using Student's t-test. (** P < 0.01, **** P < 0.0001). *n* denotes individual mice.

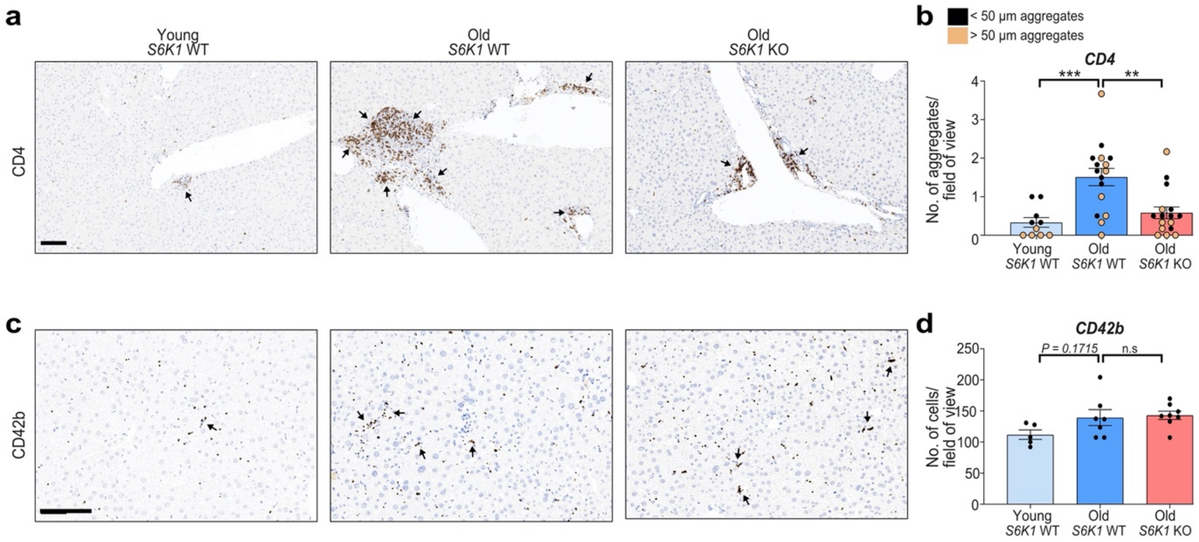


**Extended Data Figure 2. S6K1 attenuates 'inflammaging' in the liver. a,b.** CD4 staining for T-helper cells **(left)** and quantification **(right)** of livers in young *S6K1* WT (90 days; n=5), old *S6K1* WT (600 days; n=8) and old *S6K1* KO (600 days; n=8) mice. **c,d.** CD42b staining for platelets **(left)** and quantification **(right)** of livers in young *S6K1* WT (90 days; n=5), old *S6K1* WT (600 days; n=7) and old *S6K1* KO (600 days; n=8) mice. Data are expressed as mean ± SEM. Statistical significance was calculated using one-way analysis of variance with Tukey's multiple comparison test. (** P < 0.01, *** P < 0.001). *n* denotes individual mice. Scale bar, 100 μm. n.s: non-significant. WT: wild type. KO: knockout.

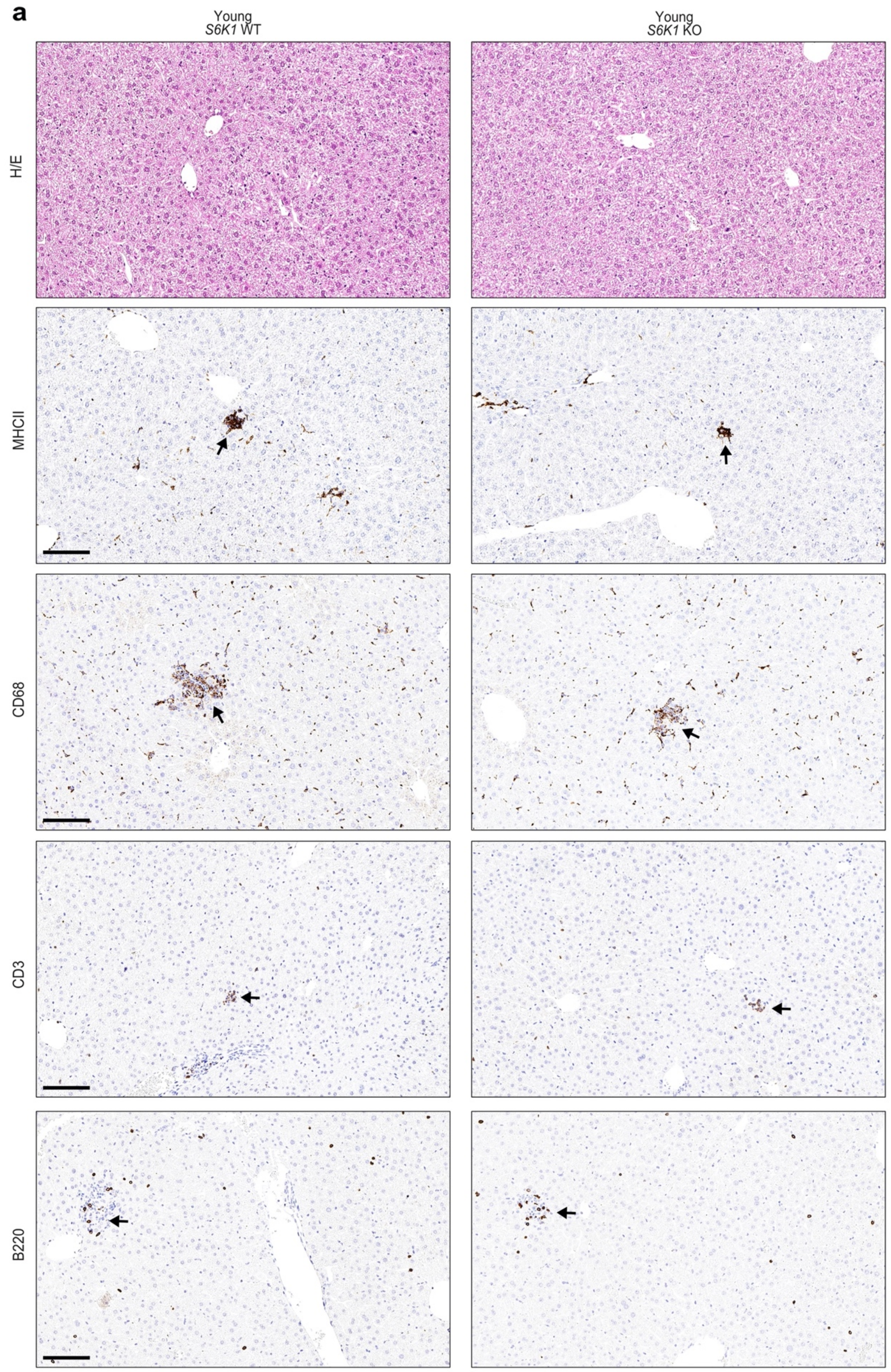


**Extended Data Figure 3. S6K1 deletion does not affect immune infiltration in young mice.** Immunohistochemistry analysis of immune cell markers in young (90

days) *S6K1* WT and KO mice. **a.** Haematoxylin and eosin (H/E) staining, MHCII staining for antigen presenting cells, CD68 staining for monocytes and macrophages, CD3 staining for t-cells and B220 staining for b-cells of livers. Scale bar, 100 μm. WT: wild type. KO: knockout.

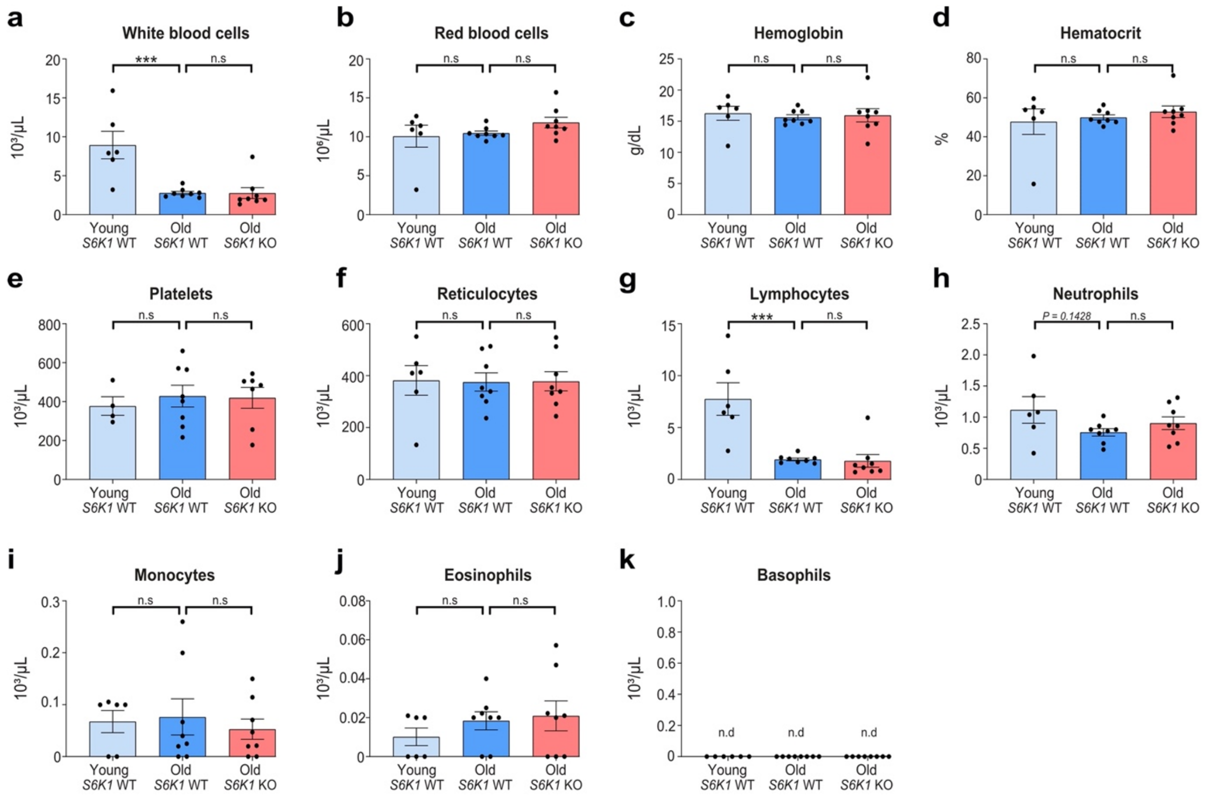


**Extended Data Figure 4. S6K1 deletion does not significantly alter the systemic blood count during aging.** Whole blood of young *S6K1* WT (90 days; n=6), old *S6K1* WT (600 days; n=8) and old *S6K1* KO (600 days; n=8) mice was used to assess the full blood count. **a.** White blood cell count ($10^3$/µL). **b.** Red blood cell count ($10^6$/µL). **c.** Haemaglobin (g/dL). **d.** Haematocrit (%). **e.** Platelet count ($10^3$/µL) in young *S6K1* WT (90 days; n=4), old *S6K1* WT (600 days; n=8) and old *S6K1* KO (600 days' n=8) mice. **f.** Reticulocytes ($10^3$/µL). **g.** Lymphocytes ($10^3$/µL). **h.** Neutrophils ($10^3$/µL). **i.** Monocytes ($10^3$/µL). **j.** Eosinophils ($10^3$/µL). **k.** Basophils ($10^3$/µL). Data are expressed as mean ± SEM. Statistical significance was calculated using one-way analysis of variance with Tukey's multiple comparison test. (* P < 0.05, ** P < 0.01, *** P < 0.001). *n* denotes individual mice. n.s: non-significant.

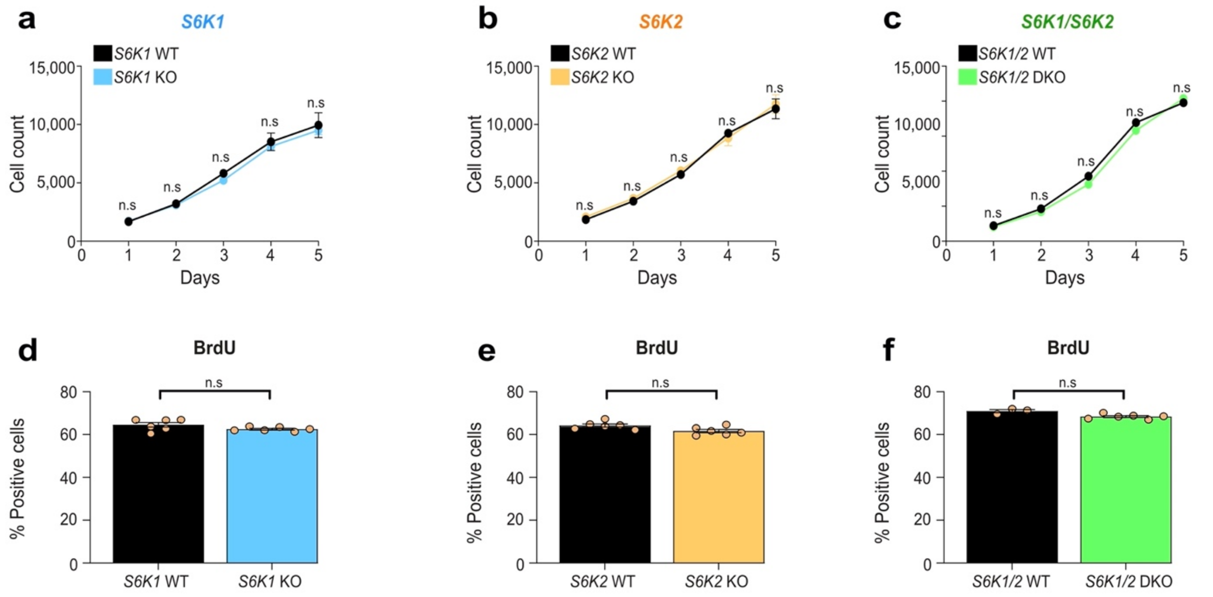


**Extended Data Figure 5. S6K1 and/or S6K2 deletion does not affect proliferation at early passage in mouse embryonic fibroblasts.** Mouse embryonic fibroblasts (MEFs) from early passage (P2) *S6K1* WT/KO, *S6K2* WT/KO and *S6K1/2* WT/DKO embryos were assessed for cell count and proliferation of the indicated genotypes. **a-c.** Time course of cell count assessed by high throughput microscopy of DAPI staining over the span of 5 days. **d-f.** Percentage of BrdU positive cells on day 1 of the indicated genotypes. Data are expressed as mean ± SEM. Statistical significance was calculated using one-way analysis of variance with Tukey's multiple comparison test. MEFs isolated from 2 independent pairs of embryos performed in triplicate wells of a single experiment. WT: wild type. KO: knockout. DKO: double knockout. n.s: non-significant. n=6 biological replicates from a single experiment. MEFs isolated from 2 independent pairs of embryos.

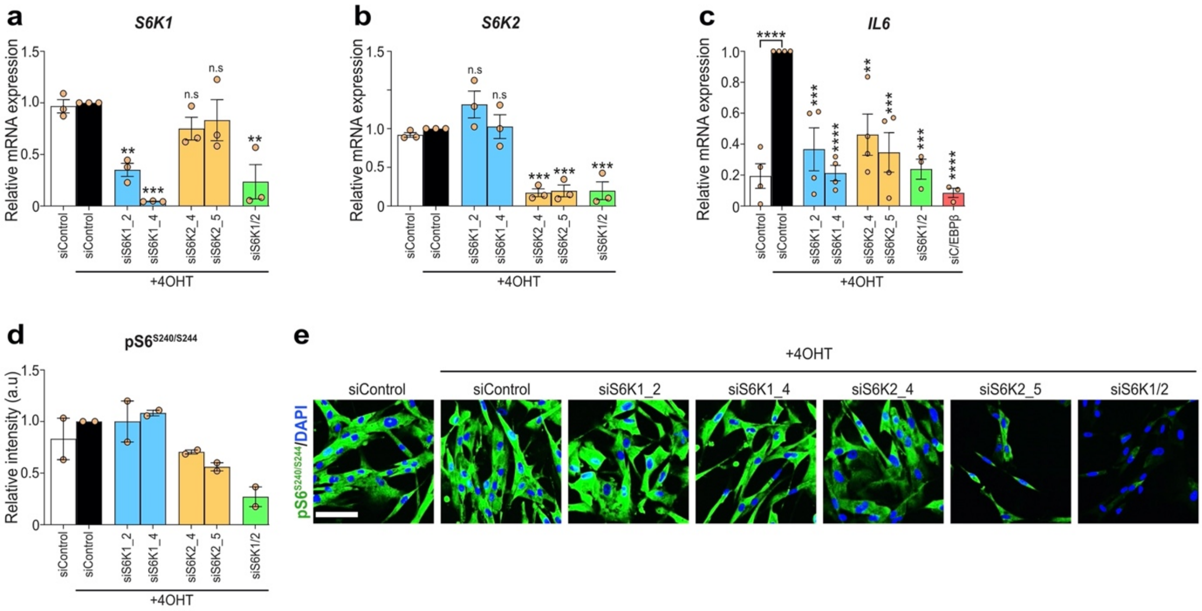

**Extended Data Figure 6. Confirmation of S6K1/2 depletion or inhibition in IMR90 ER:RAS fibroblasts.** IMR90 fibroblasts were stably transduced with the pLNC-ER: RAS retroviral vector and treated with 4-hydroxytamoxifen (4OHT) for senescence induction. **a-c.** IMR-90 ER: RAS cells were reverse transfected with either Allstars (scrambled sequence - siControl) or the indicated siRNAs. Cells were treated with or without 4OHT on the following day to induce senescence. Relative mRNA expression for *S6K1, S6K2* and *Il6* assessed by RT-qPCR- following 4 days of 4OHT treatment with the indicated siRNAs in IMR90 ER: RAS cells. *S6K1* and *S6K2:* n=3 for all conditions. *IL6:* n=4 for siControl, siS6K1_2, siS6K1_4, siS6K2_4 and siS6K2_5 and n=3 for siS6K1/2 and siC/EBPβ. mRNA expression was normalized to *Rps14* housekeeping gene. **d,e.** Quantification **(left)** and representative immunofluorescence images **(right)** for phosphorylated ribosomal protein $S6^{S240/S244}$ staining of the indicated cells (n=2). Data are expressed as mean ± SEM. Statistical significance was calculated using one-way analysis of variance with Tukey's multiple comparison test. (** $P < 0.01$, *** $P < 0.001$, **** $P < 0.0001$). n.s: non-significant. Scale bar, 100 µm.

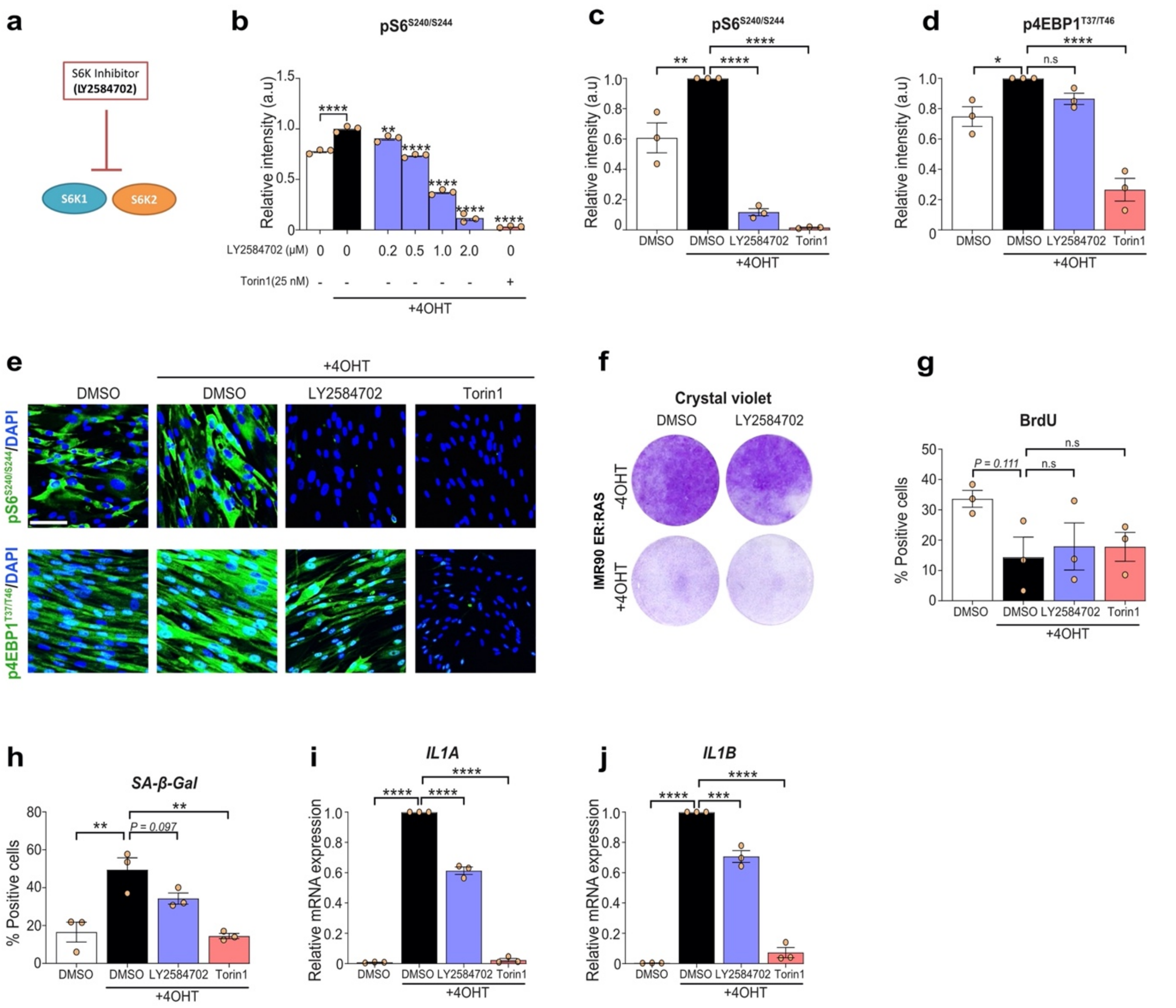


**Extended Data Figure 7. S6K1/2 inhibition in IMR90 ER:RAS fibroblasts undergoing RAS-induced senescence. a.** Schematic depicting LY2584702 inhibiting both S6K1 and S6K2. **b.** IMR90 ER: RAS cells were treated with or without 4OHT in the presence of DMSO, LY2584702 (0.2-2 μM) or Torin1 (25 nM). Quantification of immunofluorescence staining for phosphorylated ribosomal protein $S6^{S240/S244}$. n=3 biological replicates of a single experiment. **c-e.** Quantification and representative immunofluorescence images for phosphorylated ribosomal protein $S6^{S240/S244}$ and phosphorylated eukaryotic translation initiation factor 4E-binding protein 1 following 7 days of treatment with or without 4OHT in the presence of DMSO, LY2584702 (2 μM) or Torin1 (25 nM). n=3 independent experiments. **f.** IMR90 ER:

RAS cells were treated with or without 4OHT in the presence of DMSO or LY2584702 (2 μM). Cell proliferation was assessed by colony formation assay (crystal violet staining) following 13 days in culture. **g, h.** IMR90 ER: RAS cells were treated with or without 4OHT in the presence of DMSO, LY2584702 or Torin1. Quantification of IF staining for BrdU incorporation or senescence-associated beta galactosidase (SA-β-Gal) following 7 days in culture. n=3 independent experiments. **i,j.** IMR90 ER: RAS cells were treated with or without 4OHT in the presence of DMSO, LY2584702 or Torin1. Relative mRNA expression for *IL1A* and *IL1B* assessed by RT-qPCR following 6 days of 4OHT treatment. mRNA expression was normalized to *Rps14* housekeeping gene. n=3 independent experiments. Data are expressed as mean ± SEM. Statistical significance was calculated using one-way analysis of variance with Dunnett's multiple comparison test. (* $P < 0.05$, ** $P < 0.01$, *** $P < 0.001$, **** $P < 0.0001$). n.s: non-significant. Scale bar, 100 μm.

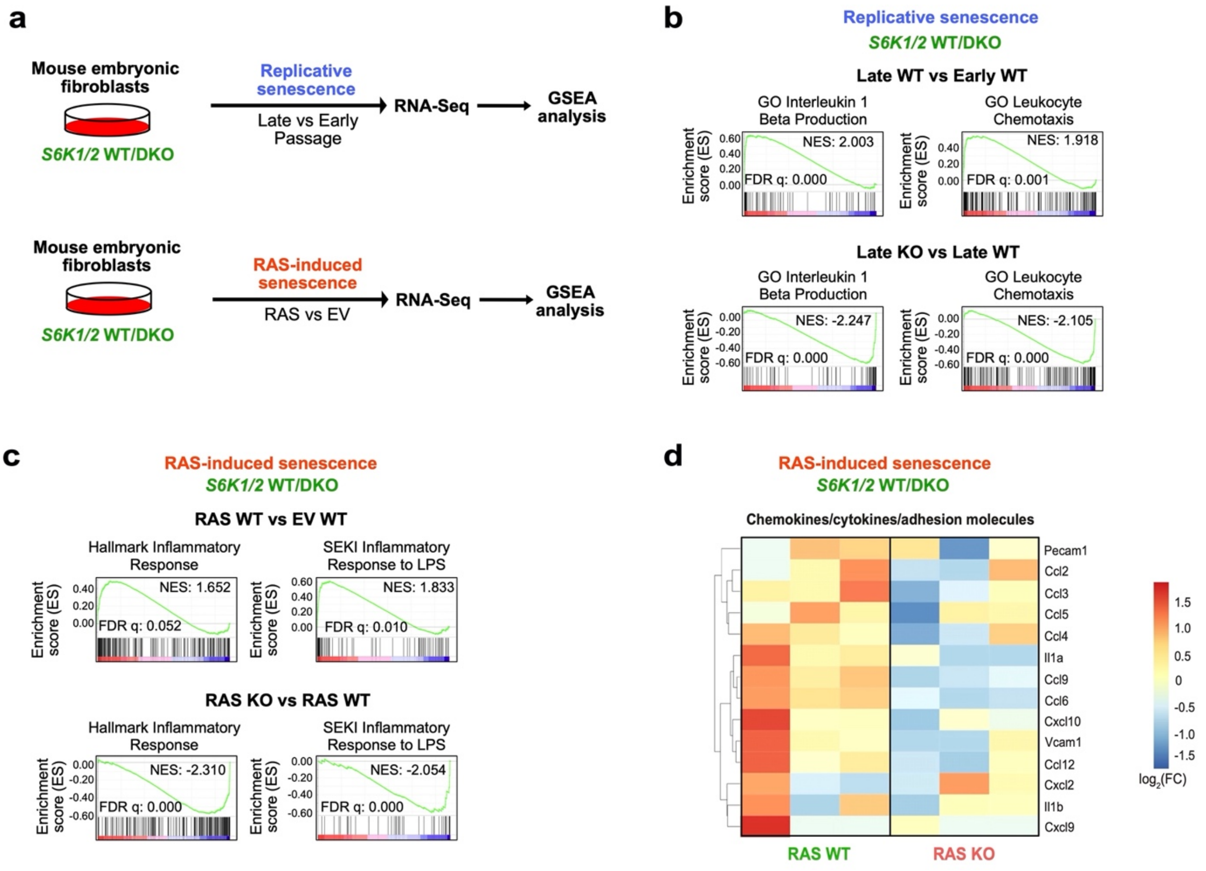


**Extended Data Figure 8. Transcriptional analysis shows that S6K1/2 regulates inflammatory pathways. a.** Experimental scheme. Mouse embryonic fibroblasts (MEFs) from S6K1/2 WT/DKO embryos were assessed for replicative senescence or RAS-induced senescence. Samples underwent subsequent RNA-sequencing and gene-set enrichment analysis (GSEA). **b.** GSEA of early S6K1/2 WT (passage 3), late S6K1/2 WT (passage 8) and late S6K1/2 DKO (passage 8) MEFs. **c.** GSEA of S6K1/2 WT MEFs expressing an empty vector (EV), S6K1/2 WT MEFs expressing $RAS^{G12V}$ or S6K1/2 DKO MEFs expressing $RAS^{G12V}$. **d.** Heatmap illustrating the gene expression pattern of key proinflammatory SASP factors involved in RAS-induced senescence. **Left:** comparison of S6K1/2 WT MEFs expressing $RAS^{G12V}$ (n=3) with S6K1/2 WT MEFs expressing EV (n=3). **Right:** comparison of S6K1/2 DKO MEFs expressing $RAS^{G12V}$ (n=3) with S6K1/2 WT MEFs expressing $RAS^{G12V}$ (n=3). NES: normalized enrichment score. FDR: false discovery rate. WT: wild type. DKO: double knockout.

### SUPPLEMENTARY TABLES

**Supplementary Table 1. List of siRNA sequences used in this study.**

| **siRNA (Human)** | **Target Sequence** |
|---|---|
| siC/EBPβ | CGGGCCCTGAGTAATCGCTTA |
| siCDKN2A ($p16^{INK4A}$) | TACCGTAAATGTCCATTTATA |
| siS6K1_2 | TCGCTTTGCATCTATCAGTAA |
| siS6K1_4 | CAGAGAGTCAATGTCATTACA |
| siS6K2_4 | TGCCATGAAAGTCCTAAGGAA |
| siS6K2_5 | ACCGCAGAGAACCGGAAGAAA |

**Supplementary Table 2. List of antibodies and dilutions used for IF and WB.**

| Target | Clone | Company | Catalogue no. | Dilution |
|---|---|---|---|---|
| S6K1 (WB) | 49D7 | Cell Signaling | #2708 | 1:1000 |
| S6K2 (WB) | Polyclonal | Cell Signaling | #14130 | 1:500 |
| Phospho-RPS6 (S240/S244) (WB/IF) | D68F8 | Cell Signaling | #5364 | 1:10,000-40,000 (WB) and 1:800 (IF) |
| Phospho-4EBP1 (T37/T46) (IF) | 236B4 | Cell Signaling | #2855 | 1:750 (IF) |
| BrdU (IF) | PRB-1 | Invitrogen | A21303 | 1:1500 |
| $p16^{INK4A}$ (IF) | JC-8 | CRUK | - | 1:750 |
| $p21^{CIP1}$ (IF) | Polyclonal | Santa Cruz | SC-471 | 1:200 |
| IL-1α (IF) | #4414 | R&D Systems | MAB200 | 1:100 |
| IL-1β (IF) | #8516 | R&D Systems | MAB201 | 1:100 |
| IL-1β (WB) | Polyclonal | Santa Cruz | SC-7884 | 1:200 |
| IL-8 (WB) | 6217 | R&D Systems | MAB208 | 1:100 |
| GAPDH (WB) | Polyclonal | Abcam | ab22555 | 1:2000 |
| HRAS (WB) | Polyclonal | Santa Cruz | SC-520 | 1:1000 |

**Supplementary Table 3. List of RT-qPCR primers used in this study**

| **Human** | **Forward primer** | **Reverse primer** |
|---|---|---|
| CCL20 | GGCGAATCAGAAGCAGCAAGCAAC | ATTGGCCAGCTGCCGTGTGAA |
| IL1A | AGTGCTGCTGAAGGAGATGCCTGA | CCCCTGCCAAGCACACCCAGTA |
| IL1B | TGCACGCTCCGGGACTCACA | CATGGAGAACACCACTTGTTGCTCC |
| IL6 | CCAGGAGCCCAGCTATGAAC | CCCAGGGAGAAGGCAACTG |
| IL8 | GAGTGGACCACACTGCGCCA | TCCACAACCCTCTGCACCCAGT |
| RPS14 | CTGCGAGTGCTGTCAGAGG | TCACCGCCCTACACATCAAACT |
| S6K1 | CGGGTACTTGGTAAAGGGGG | ATTGCCTTTTTAAGCACCTTCATGG |
| S6K2 | TTCCGGCACATGAATTGGGA | TATGTGAAGCCCAGGAAGG |

| **Mouse** | **Forward primer** | **Reverse primer** |
|---|---|---|
| *Ccl5* | CTGCTGCTTTGCCTACCTCT | CGAGTGACAAACACGACTGC |
| *Cxcl2* | CCAACCACCAGGCTACAGG | GCGTCACACTCAAGCTCTG |
| *Il1a* | CGCTTGAGTCGGCAAAGAAAT | TGGCAGAACTGTAGTCTTCGT |
| *Il1b* | TGCCACCTTTTGACAGTGATG | TGATGTGCTGCTGCGAGATT |
| $p16^{Ink4a}$ | CCCAACGCCCCGAACT | GCAGAAGAGCTGCTACGTGAA |
| $p19^{Arf}$ | TGAGGCTAGAGAGGATCTTGAGA | GCAGAAGAGCTGCTACGTGAA |
| *Rps14* | GACCAAGACCCCTGGACCT | CCCCTTTTCTTCGAGTGCTA |
